# EEG-based analysis of intrinsic brain network function in chronic pain: Insights from a comprehensive multi-data set study

**DOI:** 10.1101/2024.08.12.607584

**Authors:** Felix S. Bott, Paul Theo Zebhauser, Vanessa D. Hohn, Özgün Turgut, Elisabeth S. May, Laura Tiemann, Cristina Gil Ávila, Henrik Heitmann, Moritz M. Nickel, Melissa A. Day, Divya B. Adhia, Yoni K. Ashar, Tor D. Wager, Yelena Granovsky, David Yarnitsky, Mark P. Jensen, Joachim Gross, Markus Ploner

## Abstract

Chronic pain is associated with alterations in brain function. A better understanding of these alterations might help to develop new approaches for the diagnosis, prediction, and treatment of chronic pain. Here, we analyzed associations between chronic pain and alterations of intrinsic brain network function using resting-state electroencephalography. We included data from 537 people with chronic pain obtained from various research groups worldwide. We found strong evidence for associations between pain intensity and intrinsic brain network connectivity, but the replicability of these associations in independent data was mostly low. However, a mega-analysis revealed associations of chronic pain with salience-somatomotor network connectivity at theta frequencies and with more complex patterns of intrinsic brain network connectivity. These findings provide novel insights into brain network function in chronic pain. Moreover, they highlight the need for collaborative multi-center studies, which can be guided by the present approach to promote replicability and consistency of findings.

## Introduction

Chronic pain is a multi-faceted and debilitating condition that significantly burdens individuals and society [1, 2]. Converging lines of evidence indicate that the brain plays a central role in the development and maintenance of chronic pain [2, 3]. Thus, a better understanding of the brain’s role in chronic pain might help to develop new approaches for the diagnosis, prediction, and treatment of chronic pain. To this end, we assessed how alterations in brain function, as measured by electroencephalography (EEG), relate to chronic pain.

Functional imaging studies have revealed that chronic pain is associated with changes in the function of intrinsic brain networks [4–9]. Intrinsic brain networks are spatially extended networks of brain regions that share similar functional properties [10, 11]. Chronic pain has been mainly linked to the function of the somatomotor (SMN; aka pericentral), frontoparietal (FPN; aka lateral frontoparietal / control / central executive), salience (SN; aka midcingulo-insular / ventral attention), and default (DN; aka medial frontoparietal) networks. For instance, evidence suggests reduced connectivity between the default and salience networks and increased connectivity between the somatomotor and both the salience and frontoparietal networks [4–9]. These findings are based on hemodynamic signals obtained from functional magnetic resonance imaging (fMRI). Neurophysiological recordings of brain activity using electroencephalography (EEG) can complement these findings by assessing intrinsic brain network function at shorter time scales. Moreover, EEG is broadly available, cost-efficient, and potentially mobile, thus facilitating the translation of findings to clinical practice. Hence, EEG has been increasingly used to investigate intrinsic brain network function in neuropsychiatric disorders [12, 13]. However, EEG has not been used to investigate intrinsic brain network function in chronic pain so far.

In the present study, we used EEG to investigate the function of intrinsic brain networks in people with chronic pain. We assessed the replicability and consistency of findings based on seven independent data sets (n = 537) including two data sets recorded by our research group in Germany and five data sets recorded by research groups in Israel, New Zealand, the US, and Australia. We focused on connectivity across intrinsic brain networks, as connectivity-based assessments have proven particularly informative in various neuropsychiatric disorders, including chronic pain [13–17]. We linked brain network connectivity to self-reported pain intensity of people with chronic pain using both univariate correlation and multivariate regression techniques.

We specifically addressed the following research questions (Fig. 1). First, we assessed how pain intensity relates to connectivity in intrinsic brain networks (*network analysis*). Second, we assessed how pain intensity relates to standard EEG features previously linked to chronic pain (*standard analysis*). Third, as a positive control analysis, we assessed how age relates to intrinsic brain network connectivity and standard EEG features (*control analysis*). Fourth, we explored how multivariate models using brain connectivity beyond the a-priori-defined intrinsic brain networks could predict pain intensity and analyzed which aspects of the models enabled the predictions (*extended network analysis*).

**Figure 1:**
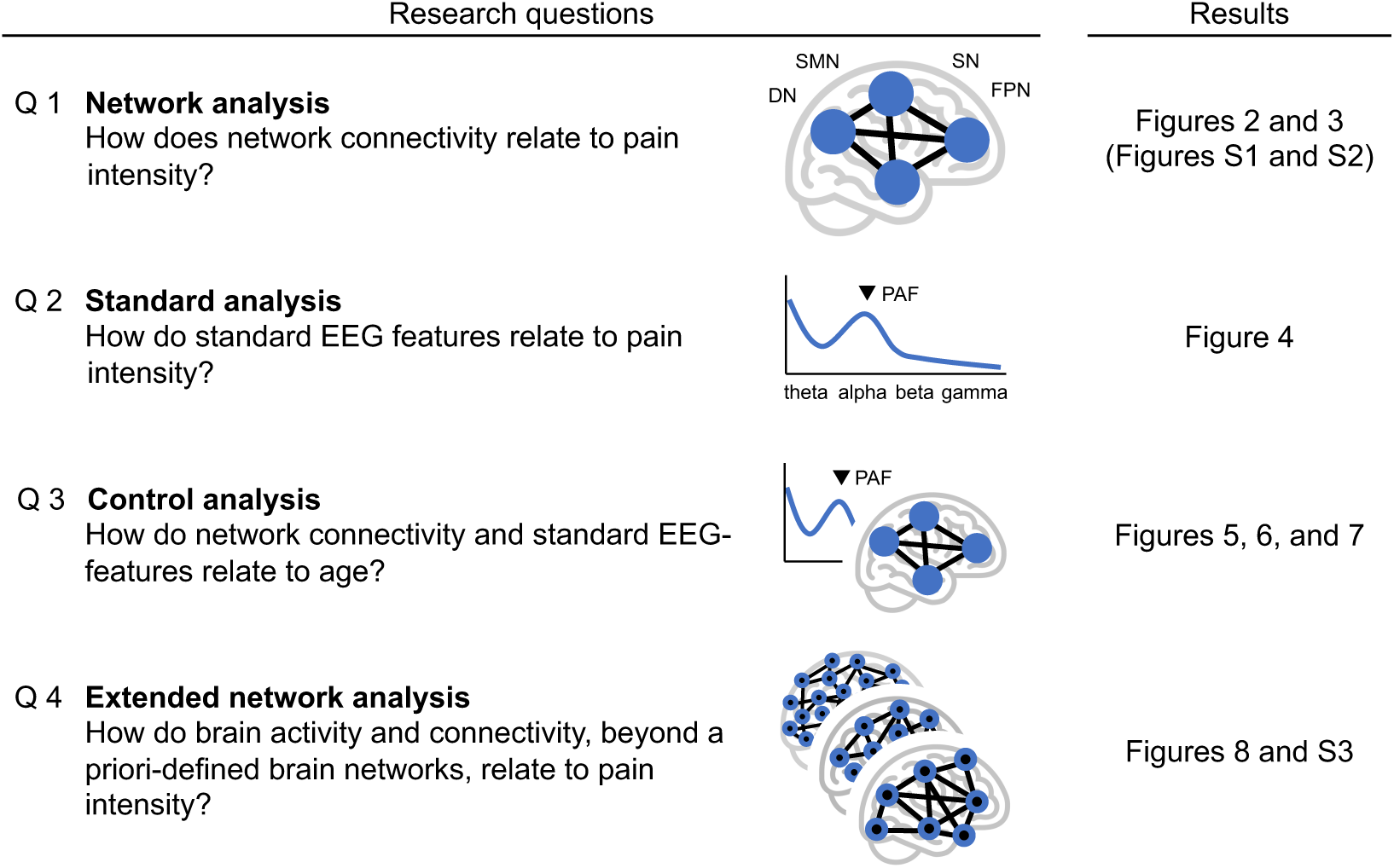
Research questions. We used EEG data from seven independent studies involving people with chronic pain to investigate how brain connectivity relates to pain intensity.

## Results

In the present study, we investigated how the connectivity between brain networks, assessed by EEG, relates to pain intensity in people with chronic pain. To this end, we analyzed seven resting-state EEG data sets of people with chronic pain (total n = 537, Table 1). The participants had different types of chronic pain, with chronic back pain being the most frequent type (n = 328). The data sets were recorded at various sites with different devices and setups but were centrally collected and analyzed at one site. To assess the relationship between brain network connectivity and pain intensity and the replicability of the findings, we pursued a discovery-replication approach. The discovery data set was a large data set recorded by our group in Munich (n = 119). The replication data sets were six other data sets recorded by our group in Munich and at other sites (n = 47 to 123, total n = 418). In addition, we performed mega-analyses by analyzing the joint data set comprising all seven data sets (n = 537).

**Table 1:**
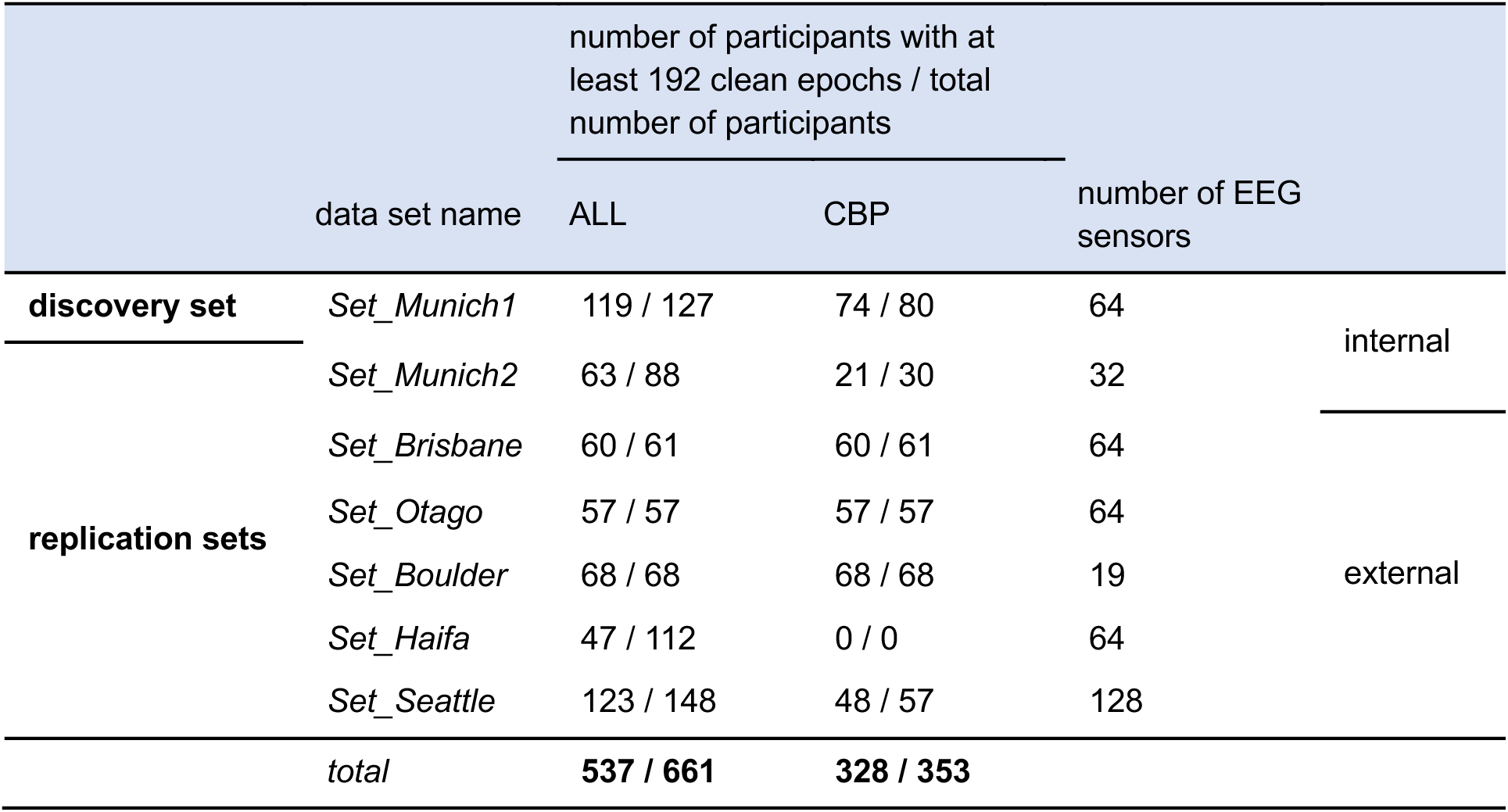
Overview of data sets. The study included seven data sets, each comprising resting-state EEG and metadata of people with different types of chronic pain (ALL). Chronic back pain (CBP) was the most frequent type of chronic pain. To ensure accurate EEG feature estimates, we included only participants with a minimum number of 192 clean EEG data epochs (see methods section for details). The numbers preceding and following the slash in columns ALL and CBP represent the counts of participants with at least 192 clean epochs and the total number of participants, respectively.

We performed univariate analyses to assess associations between connectivity of individual brain network pairs and pain intensity. In addition, we performed multivariate analyses to investigate associations between patterns of brain network connectivity and pain intensity. Univariate and multivariate analyses relied on Bayesian statistics, which allow for interpreting both positive and negative findings. We interpreted Bayes factors (BF) > 1, > 3, and > 10 (<1, <1/3, <1/10) as anecdotal, moderate, and strong evidence in favor of (or against) an effect.

We first analyzed the discovery data set and then used the replication sets to quantify the replicability and consistency of the effects. *Replicability* of effects in independent data was quantified by a BF reflecting the strength of evidence for (or against) the same correlation between EEG features and pain intensity in the pooled replication sets. Evidence for (against) replicability was interpreted as anecdotal, moderate, or strong, as defined above. To assess the *consistency* of effects, we developed a score quantifying how associations between EEG features and pain intensity could be replicated in the individual replication sets (see supplementary materials). This consistency score is directly proportional to the sum of (1) the number of replication sets for which the direction of the correlation equals that in the discovery set and (2) the number of replication sets in which, in addition, the correlations yield a BF > 3. It has four levels, reflecting no, anecdotal, moderate, and strong evidence for consistency. Under the Null hypothesis, the probabilities of observing at least anecdotal, moderate, or strong evidence for consistency are p < 0.16, < 0.05, and < 0.01, respectively. Finally, we performed mega-analyses on the joint data set comprising the discovery and all replication sets.

The study was preregistered at osf.io (https://osf.io/4qmyw/).

### Network analysis: How does brain network connectivity relate to pain intensity?

The network analysis focused on connectivity between four brain networks that have previously been linked to neuropsychiatric disorders and chronic pain [4, 18]: the frontoparietal (FPN), the salience ventral attention (SN), the default (DN), and the somatomotor network (SMN). To spatially define these brain networks, we employed the Yeo atlas [10]. To evaluate connectivity between brain networks, we extracted signals representative of the networks and computed the amplitude envelope correlation (AEC, [19]) between these signals. We used an amplitude-based connectivity metric as it is conceptually close to fMRI-based connectivity assessments on which the intrinsic brain network concept is based [10]. Connectivity values were extracted for three frequency bands that figure prominently in network communication: theta (3-8 Hz), alpha (8-13 Hz), and beta (13-30 Hz). This resulted in 18 connectivity values for each participant. We related these connectivity values to pain intensity as measured by ratings of the average pain intensity over the past one to four weeks (depending on the data set).

#### Univariate analyses

First, we sought to determine how the connectivity of each brain network pair relates to pain intensity in the discovery set (Fig. 2a). Pain intensity positively correlated with all connectivity values at theta frequencies and negatively correlated with all connectivity values at alpha frequencies. Evidence for a correlation was moderate to strong for three network pairs in the theta (FPN-SN: r = 0.21, BF = 3.7; DN-SN: r = 0.26, BF = 18; DN-FPN: r = 0.25, BF = 14) and for one network pair in the alpha band (FPN-SN: r = −0.33, BF = 254). We found moderate evidence against correlations in the beta band.

**Figure 2:**
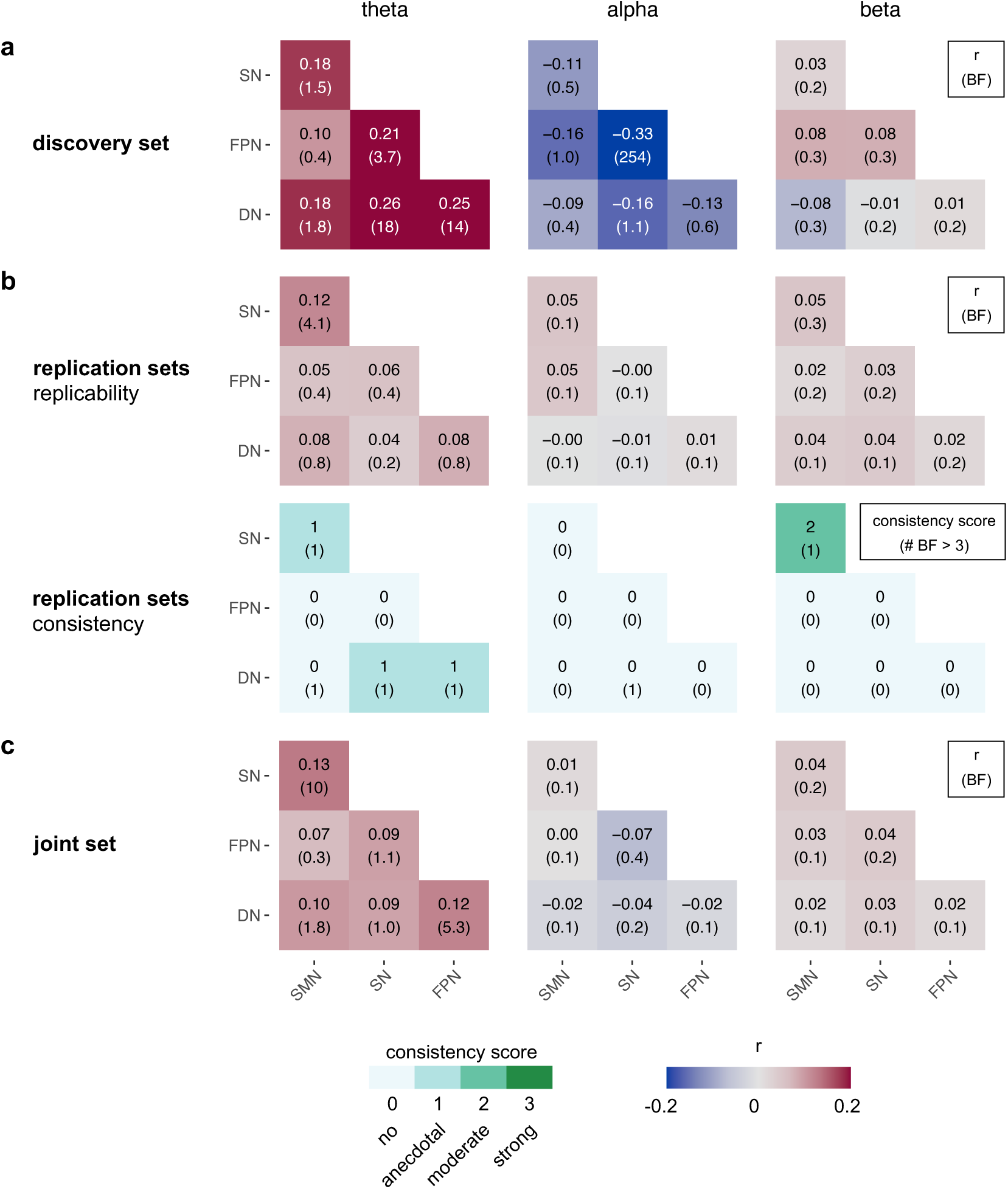
Univariate correlations between brain network connectivity at theta, alpha, and beta frequencies and pain intensity. (a) Correlations between pain intensity and brain network connectivity in the discovery set. Each heatmap tile’s top number and color represent the correlation coefficient; the bottom number is the associated BF. (b) Upper panel row: Correlations in the pooled replication sets. The meanings of numbers and colors are match those of panel (a). Lower panel row: Summary of correlations observed in the replication sets. The top number and color of each heatmap tile represent the consistency score. The bottom number represents the number of replication data sets that correlate with BF > 3 in the same direction as in the discovery set. (c) Correlations in the joint set. The meanings of numbers and colors match those of panel (a). SMN, somatormotor network; SN, salience network; FPN, frontoparietal network; DN, default network.

Second, we assessed the replicability and consistency of the effects in the six replication data sets (Fig. 2b). The results revealed that both replicability and consistency were low. Only for the SMN-SN connection in the theta band, which had yielded anecdotal evidence for an effect in the discovery set, we found moderate evidence for replicability and anecdotal evidence for consistency.

Third, we performed a mega-analysis on the joint data set (Fig. 2c). The results provided strong evidence for a positive correlation between SMN-SN connectivity at theta frequencies and pain intensity (r=0.13, BF = 10.5). Also at theta frequencies, there was moderate evidence for a positive correlation between DN-FPN connectivity and pain intensity (r = 0.12, BF = 5.3). Moreover, there was mostly inconclusive evidence for the remaining connections at theta frequencies and evidence against correlations at alpha and beta frequencies.

#### Multivariate analyses

Next, we investigated how patterns of brain network connectivity relate to pain intensity. To this end, we trained and tested different machine learning (ML) models that employ connectivity values at theta, alpha, and beta frequencies as features. A model trained and tested on the discovery data set yielded strong evidence for a cross-validated correlation between predicted and observed pain intensity (“in-sample” cross validation (CV), r = 0.35, BF = 688, permutation-based p < 0.001, Fig. 3a and b).

**Figure 3:**
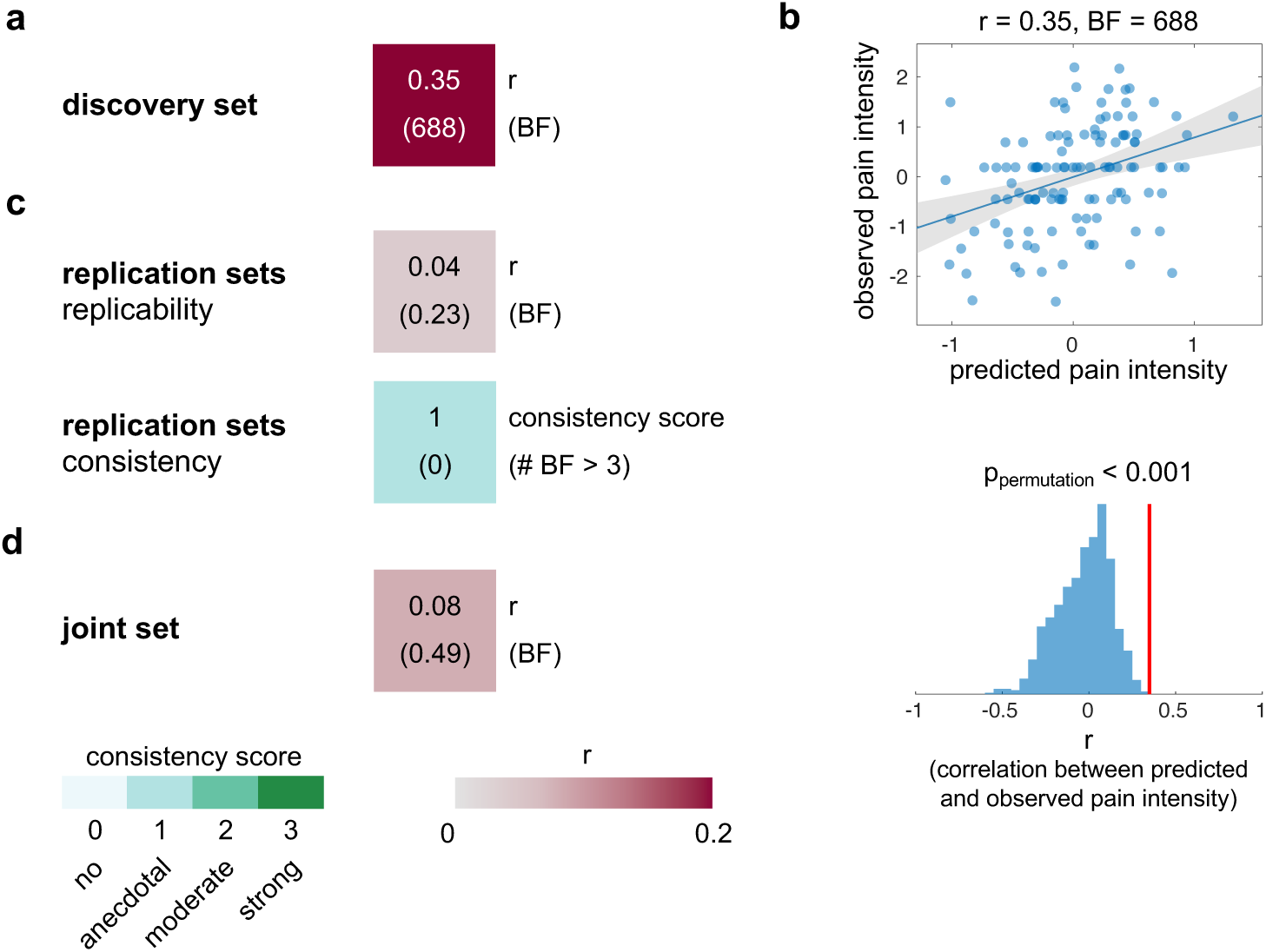
Associations between multivariate patterns of brain network connectivity and pain intensity. (a) In-sample, leave-one-participant-out cross-validated (LOO-CV) correlation between predicted and observed pain intensity in the discovery set. (b) Scatter plot of the cross-validated correlation between predicted and observed pain intensity and the result of a permutation test, confirming the statistical significance of predictions. (c) Upper tile: Out-of-sample correlations between predicted and observed pain intensity in the pooled replication sets. The meanings of numbers and colors match those in panel (a). Lower tile: Summary of out-of-sample correlations observed in individual replication sets. The top number and color of each tile represent the consistency score. The bottom number represents the number of replication data sets that exhibit a positive correlation with BF > 3. (d) In-sample, cross-validated (LOO-CV) correlation between predicted and observed pain intensity in the joint set. The meanings of numbers and colors match those in panel (a).

We then tested the model in the six replication sets (“out-of-sample” validation, Fig. 3c). The results revealed that its prediction performance did not generalize to independent data. We found evidence against a correlation between predicted and observed pain intensity (r = 0.04, BF = 0.23) and only anecdotal evidence for consistency in the replication sets.

In the multivariate mega-analysis (Fig. 3d), i.e., when training and testing a model on the joint set, no connectivity pattern related to pain intensity (r = 0.08, BF = 0.49) was detected.

#### Subgroup analysis

To investigate how brain network connectivity relates to pain intensity in more homogenous groups of people with chronic pain, we performed a subgroup analysis in the largest subgroup of study participants, i.e., those with chronic back pain (CBP) (Fig. S1 and S2). In this subgroup, the pattern of univariate correlations between brain network connectivity and pain intensity was qualitatively similar to that of all people with chronic pain, but replicability and consistency were low. ML models trained and tested on the CBP subgroup of the discovery or the joint set did not detect patterns of brain network connectivity related to pain intensity.

#### Summary

Univariate analysis of the discovery set showed positive correlations between brain network connectivity at theta frequencies and pain intensity, and negative correlations between connectivity at alpha frequencies and pain intensity. Multivariate analysis of the discovery set revealed a connectivity pattern related to pain intensity. However, the replicability and consistency of results were low. The mega-analysis of the combined data sets revealed a positive correlation between SMN-SN connectivity at theta frequencies and pain intensity.

### Standard analysis: How do standard EEG features relate to pain intensity?

We next assessed how standard EEG features other than brain network connectivity relate to pain intensity. To this end, we assessed global signal power at theta, alpha, beta, and gamma frequencies and the peak alpha frequency (PAF), as these features have been most frequently investigated in chronic pain [20]. Considering previous evidence [20], we conducted one-sided tests to determine whether theta and beta power positively correlated and PAF negatively correlated with pain intensity. As no hypotheses pertaining to directionality existed for alpha and gamma power, we assessed these correlations to pain intensity bidirectionally. We found moderate evidence against a correlation with pain intensity for all features (BF < 1/3 for all EEG features in the discovery set (Fig. 4a), BF < 1/3 for all EEG features in the replication sets (Fig. 4b)). Likewise, the mega-analysis yielded evidence against correlations between EEG features and pain intensity (Fig. 4c). Together, the standard analysis yielded evidence against correlations between commonly analyzed EEG features and pain intensity.

**Figure 4:**
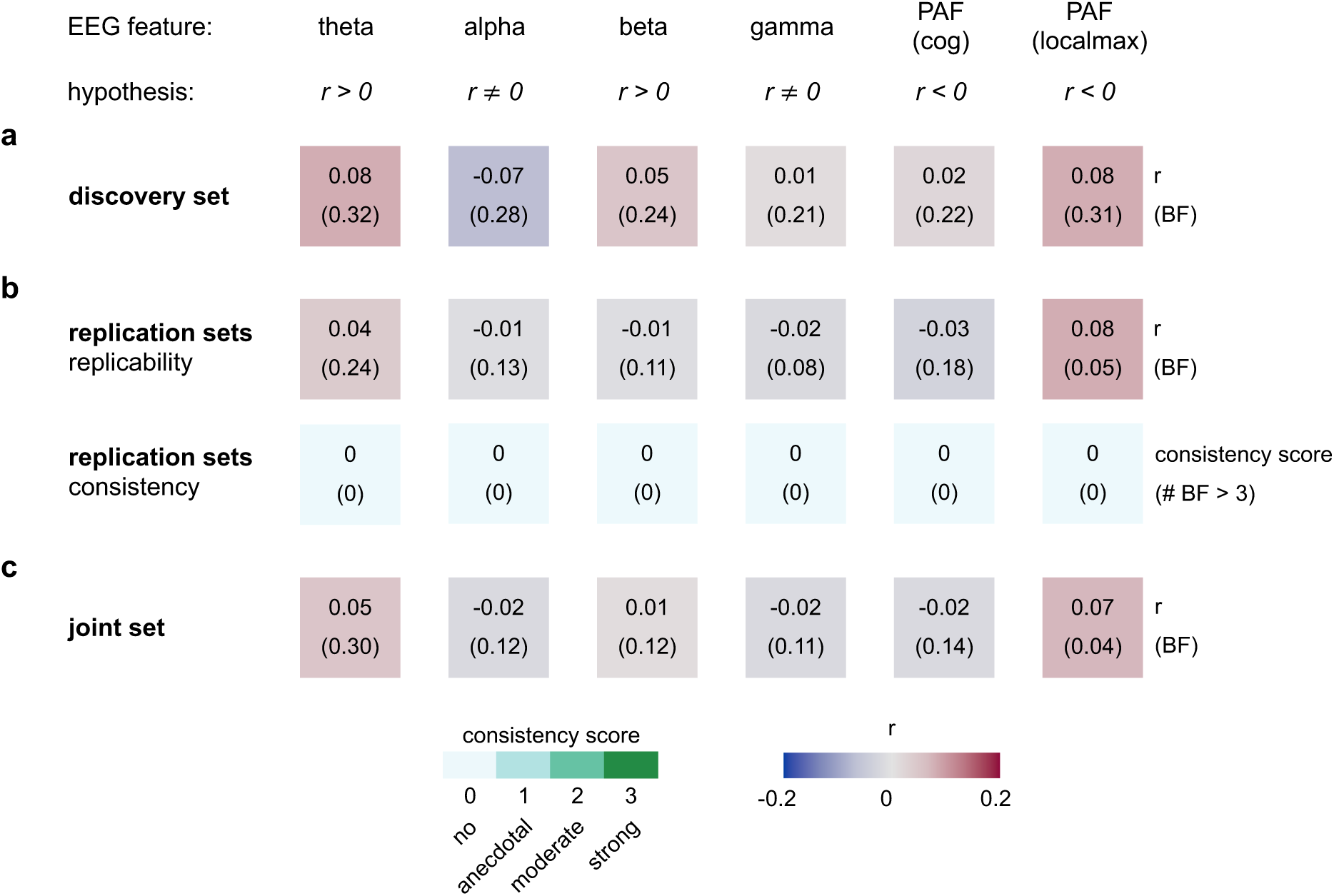
Univariate correlations between standard EEG features and pain intensity. (a) Correlations between pain intensity and EEG features in the discovery set. Each tile’s top number and color represent the correlation coefficient, and the bottom number is the associated BF. (b) Top tile row: Correlations in the pooled replication sets. The meanings of numbers and colors are match those in panel (a). Bottom tile row: Summary of correlations observed in the replication sets. The top number and color of each tile represent the consistency score. The bottom number represents the number of replication data sets that correlate with BF > 3 in the same direction as in the discovery set. (c) Correlations in the joint set. The meanings of numbers and colors match those of panel (a).

### Control analysis: How do brain network connectivity and standard EEG features relate to age?

Next, we aimed to check the data quality and the methodology’s sensitivity by analyzing how the EEG features relate to age, which is known to affect many measures of brain function [21–23]. Univariate analyses revealed negative correlations between brain network connectivity and age at all frequencies, with the strongest correlations in the theta band (Fig. 5a). The frequency specificity of effects was reflected in the replication and consistency patterns (Fig. 5b). We found moderate to strong evidence for replicability in all six connections at theta frequencies and evidence against replicability in all connections at alpha and beta frequencies. Moreover, there was anecdotal to strong evidence for consistency at theta frequencies and mostly no evidence for consistency at alpha and beta frequencies. The mega-analysis of the joint set yielded strong evidence for a negative correlation of theta band connectivity of all network pairs with age (Fig. 5c). Mostly evidence against correlations was found in the alpha and beta frequency bands.

**Figure 5:**
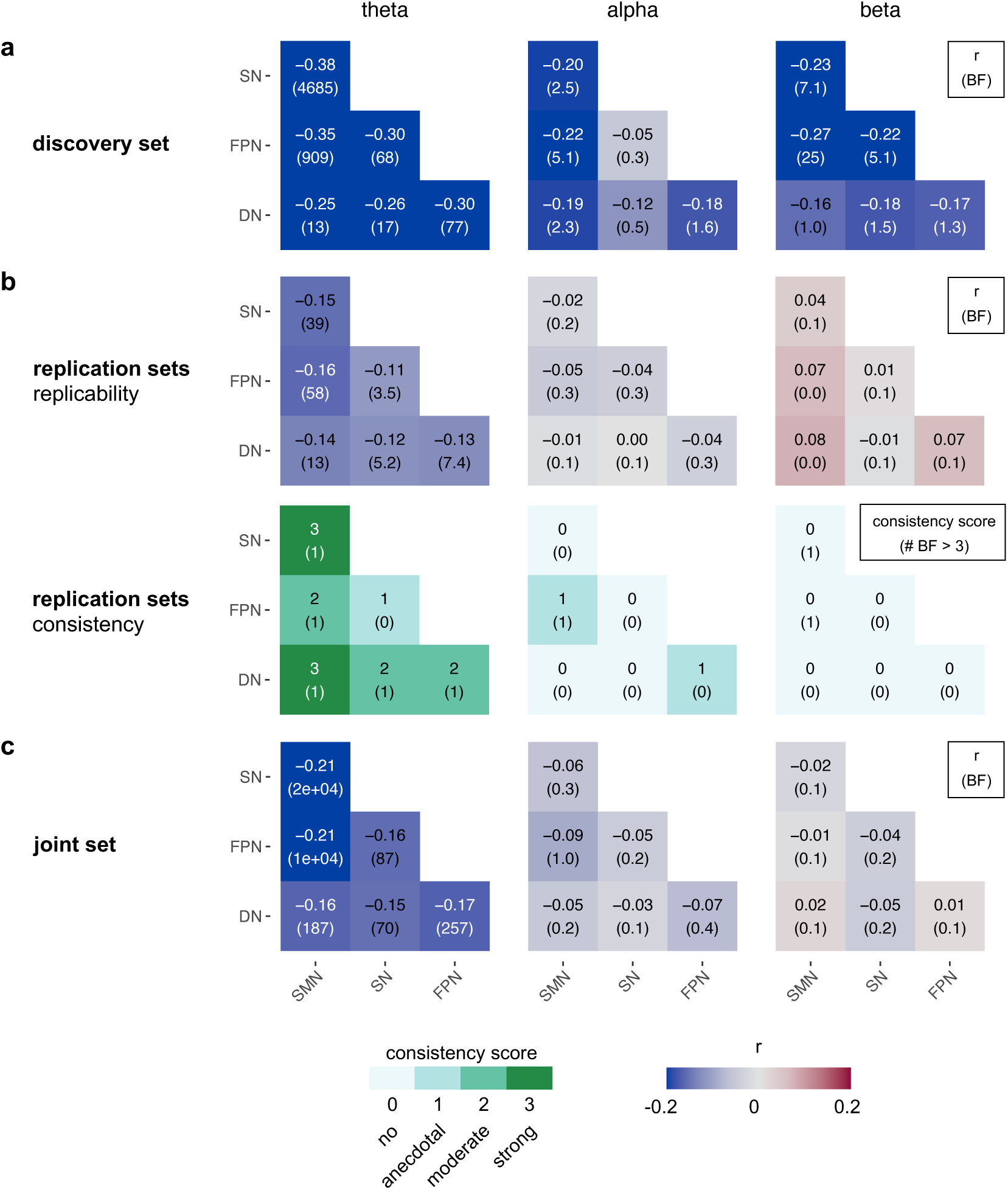
Univariate correlations between brain network connectivity at theta, alpha, and beta frequencies and age. (a) Correlations in the discovery set. Each heatmap tile’s top number and color represent the correlation coefficient, the bottom number the associated BF. (b) Top panel row: Correlations in the pooled replication set. The meanings of numbers and colors match those of panel (a). Bottom panel row: Summary of correlations observed in the replication sets. The top number and color of each heatmap tile represent the consistency score. The bottom number represents the number of replication data sets that exhibit a correlation with BF > 3 in the same direction as in the discovery set. (c) Correlations in the joint set. The meanings of numbers and colors match those of panel (a). SMN, somatormotor network; SN, salience network; FPN, frontoparietal network; DN, default network.

Multivariate analyses showed that an ML model trained and tested on the discovery data set predicted age significantly better than chance (Fig. 6a and b, r = 0.41, BF > 10^3^, permutation-based p < 0.001). We found some evidence that the model generalized to independent data sets (Fig. 6c). There was moderate evidence for the replicability (r = 0.12, BF = 6.23) and strong evidence for the consistency of model performance. In the multivariate mega-analysis of the joint set, the ML algorithm yielded strong evidence for a relationship between brain network connectivity patterns and age (Fig. 6d, r = 0.22, BF > 10^3^).

**Figure 6:**
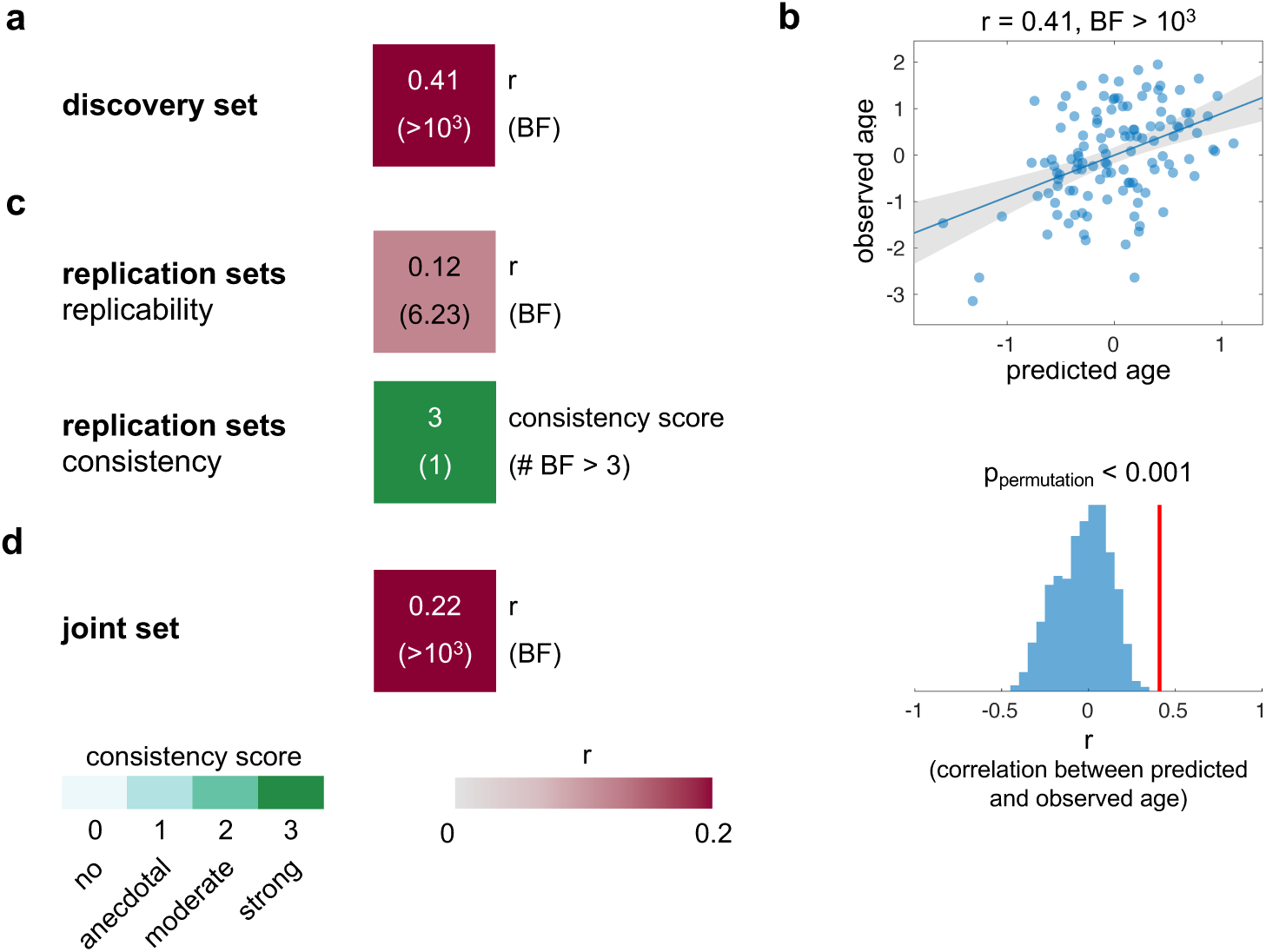
Associations between multivariate patterns of brain network connectivity and age. (a) In-sample, cross-validated (LOO-CV) correlation between predicted and observed age in the discovery set. (b) Scatter plot of the cross-validated correlation between predicted and observed age and the result of a permutation test, confirming the statistical significance of predictions. (c) Top tile: Out-of-sample correlations between predicted and observed age in the pooled replication sets. The meanings of numbers and colors match those in panel (a). Bottom tile: Summary of out-of-sample correlations observed in individual replication sets. The top number and color of each tile represent the consistency score. The bottom number represents the number of replication data sets that exhibit a positive correlation with BF > 3. (d) In-sample, cross-validated (LOO-CV) correlation between predicted and observed age in the joint set. The meanings of numbers and colors match those in panel (a).

We further assessed the correlations between standard EEG features and age (Fig. 7). We mostly found evidence against a correlation between global signal power at theta, alpha, beta, and gamma frequencies and age (Fig. 7a to 7c). However, in the discovery set, we found strong evidence for a negative correlation between age and PAF as quantified by two different methods (center of gravity (cog), local maximum (localmax)) (Fig. 7a). For both methods, there was strong evidence for both consistency and replicability of effects (Fig. 7b). Lastly, the mega-analysis of the joint set yielded strong evidence for a negative correlation between age and PAF (Fig. 7c) and evidence against correlations between age and other standard EEG features.

**Figure 7:**
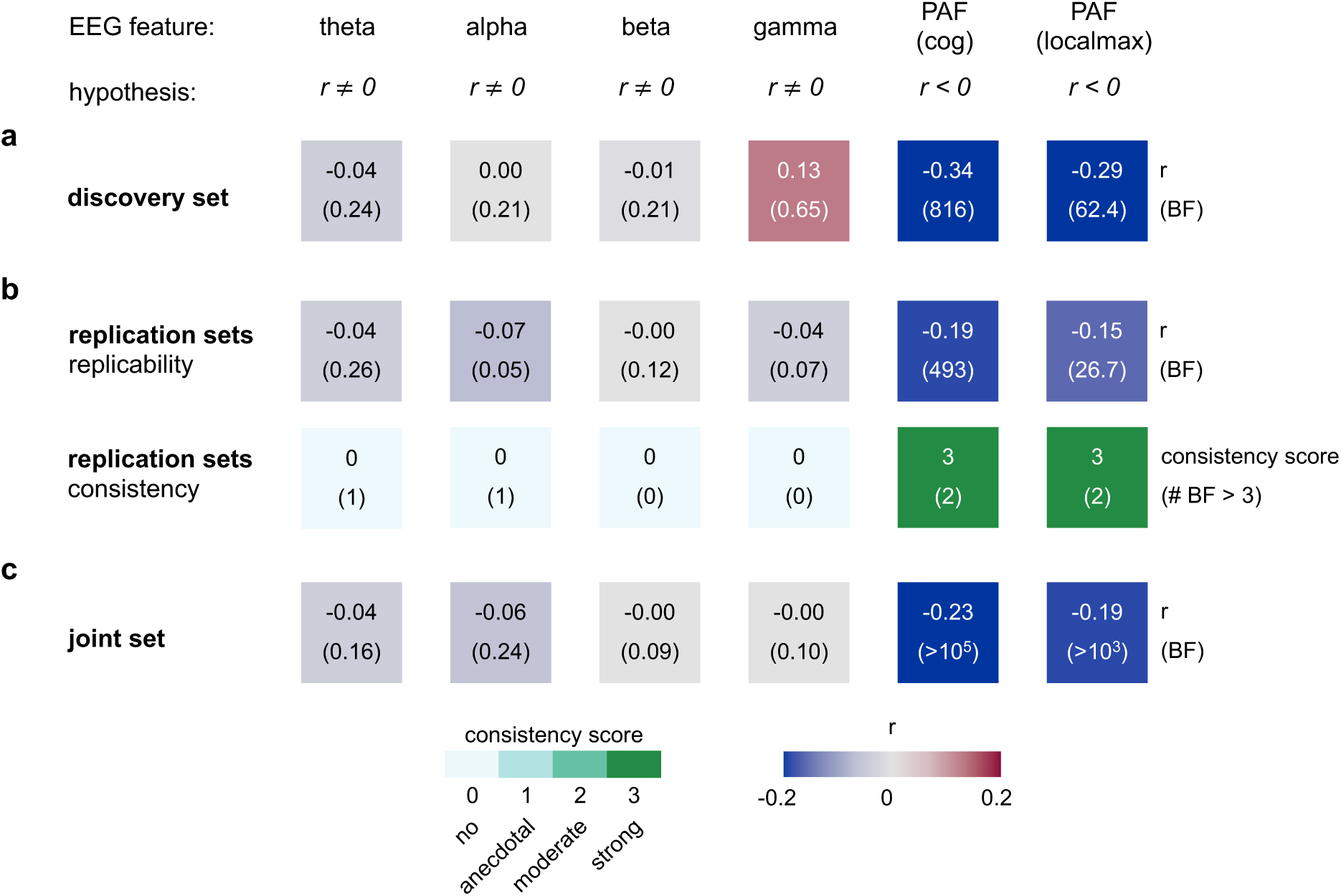
Univariate correlations between standard EEG features and age. (a) Correlations in the discovery set. Each tile’s top number and color represent the correlation coefficient, and the bottom number is the associated BF. (b) Top tile row: Correlations in the pooled replication sets. The meanings of numbers and colors match those of panel (a). Bottom tile row: Summary of correlations observed in the replication sets. The top number and color of each tile represent the consistency score. The bottom number represents the number of replication data sets that exhibit a correlation with BF > 3 in the same direction as in the discovery set. (c) Correlations in the joint set. The meanings of numbers and colors match those of panel (a).

In summary, the control analysis yielded strong and replicable evidence that univariate brain network connectivity at theta frequencies, multivariate patterns of network connectivity across all frequencies, and peak alpha frequency relate to age.

### Extended network analysis: How do brain activity and connectivity, beyond a priori-defined brain networks, relate to pain intensity?

In our primary analysis, we assessed connectivity across four a-priori-defined intrinsic brain networks using an a-priori-defined approach to compute connectivity. However, other approaches to brain network connectivity might be equally or more informative about pain intensity. We, therefore, varied the spatial definition of brain networks and the approach to computing connectivity. Specifically, we assessed connectivity among all seven intrinsic brain networks proposed by Yeo (Yeo-7), among 25 anatomically defined brain regions, and among 100 uniformly distributed brain locations. Moreover, we varied the approaches to extract representative signals. In addition to global orthogonalization, we considered pairwise orthogonalization and standard PCA to compute representative signals. Furthermore, in addition to computing the AEC based on the logarithm of the squared signal amplitudes (as proposed in [19]), we computed it directly based on the signal envelopes (as done in [24]). All these conceptual and methodological variations were integrated as feature candidates in a single extended ML model (see methods and supplementary materials for details).

The results are displayed in Fig. 8. The first column of Fig. 8a shows that the extended ML model achieved a higher prediction accuracy in the discovery data set compared to the model of the primary analysis (prediction performance of ML models from primary analysis are reiterated in the second column of Fig. 8a). Moreover, the extended model showed anecdotal evidence for the replicability (r = 0.11, BF = 2.34) and moderate evidence for the consistency of prediction performance across replication sets. In addition, when training and testing the model on the joint set, the algorithm yielded strong evidence for a brain connectivity pattern related to pain intensity (r = 0.19, BF > 10^3^). Thus, other approaches to brain connectivity than our primary analysis appear to be more informative about pain intensity.

**Figure 8:**
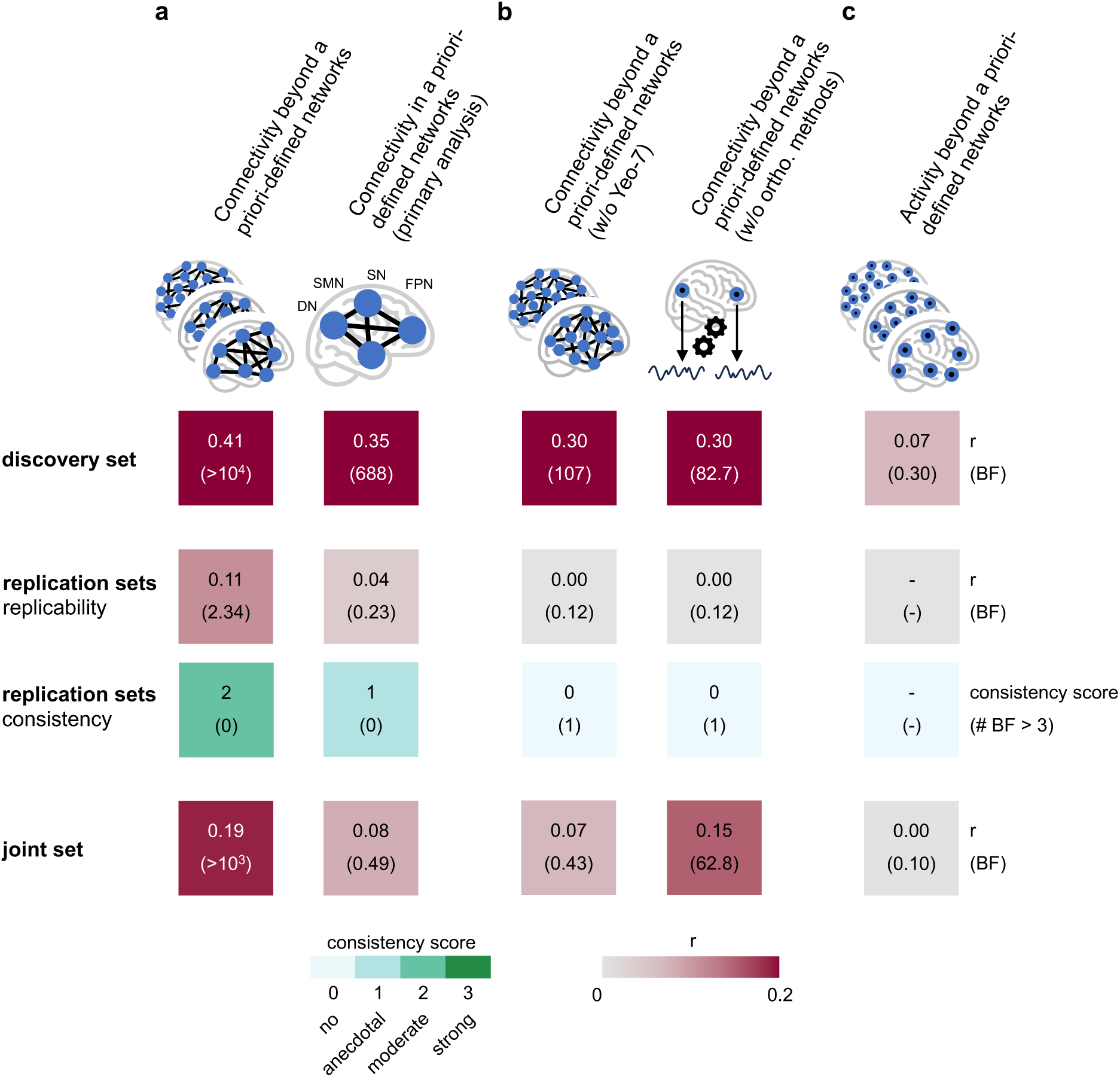
Associations between multivariate patterns beyond the four a priori-defined brain networks and pain intensity. (a) The first column summarizes the performance of the extended ML model which is based on an extended feature space comprising different network and connectivity definitions. The second column repeats the performance of the ML model of the primary analysis. Panels (b) and (c) summarize the performances of variants of the extended ML model. (b) The first column pertains to a variant in which the feature space does not comprise measures of connectivity estimates based on the Yeo-7 intrinsic brain networks. The second column pertains to a variant in which solely connectivity features were considered that did not rely on the proposed orthogonalization approaches to compute representative signals. (c) Summary of the performance of a variant of the extended ML model which uses measures of brain activity as features instead of measures of brain connectivity. (a) – (c) First tile row (discovery set): In-sample, cross-validated (100-fold-CV) correlation between predicted and observed pain intensity in the discovery set. Second tile row (replication sets – replicability): Out-of-sample correlations between predicted and observed pain intensity in the pooled replication sets. The meanings of numbers and colors match those of the first tile row. Third tile row (replication sets – consistency): Summary of out-of-sample correlations observed in individual replication sets. The top number and color of each tile represent the consistency score. The bottom number represents the number of replication data sets that exhibit a positive correlation with BF > 3. Fourth tile row: In-sample, cross-validated (100-fold-CV) correlation between predicted and observed pain intensity in the joint set. The meanings of numbers and colors match those of the first tile row.

Next, we aimed to understand which approach to brain connectivity is most informative. In all folds of the 100-fold-CV, in the discovery and the joint set, the models utilized connectivity estimates based on the seven intrinsic brain networks defined by Yeo and computed via one of the proposed orthogonalization methods for representative signals. To confirm that these connectivity estimates are more informative than other variants, we performed ablation analyses (Fig. 8b). We trained and tested extended models and omitted connectivity based on the Yeo-7 intrinsic brain networks from the set of candidate features. The prediction accuracies of the resulting models were substantially worse than those of the full models, indicating that connectivity among the Yeo-7 intrinsic brain networks is more informative. We further trained and tested extended models excluding from the set of candidate features all connectivity estimates computed via one of the proposed orthogonalization methods for representative signals. The resulting prediction accuracies were below those of the full model, indicating that the proposed orthogonalization methodology yields connectivity estimates that can be more informative about pain intensity than connectivity estimates computed via standard PCA.

Finally, we investigated whether neural connectivity is more informative about pain intensity than regional brain activity. We, therefore, compared the performance of extended models, which employ brain connectivity as features, to models that employ brain activity as features (Fig. 8c). The in-sample prediction accuracies of the activity-based models were below chance level (discovery set: r = 0.07, BF = 0.30; joint set: r = 0.0, BF = 0.10). These findings confirm that brain connectivity is more informative about chronic pain intensity than brain activity.

## Discussion

Understanding and objectifying the brain mechanisms of chronic pain is a key challenge in pain research. In the present study, we therefore investigated how chronic pain intensity relates to EEG measures of brain connectivity. We scrutinized the replicability and consistency of these associations by analyzing seven independent data sets, resulting in the largest analysis of EEG data in people with chronic pain so far. Considering converging evidence from fMRI studies, we focused on analyzing connectivity between intrinsic brain networks. These *network analyses* showed some associations between pain intensity and connectivity, but the consistency and replicability were low. However, there were indications of a relationship between pain intensity and SMN-SN connectivity at theta frequencies, as well as between pain intensity and connectivity patterns based on all seven intrinsic brain networks. Thus, assessing these features in future collaborative multi-center studies guided by the present approach is a promising way to gain new insights into the brain mechanisms of chronic pain.

### Network analysis

In the discovery data set, we found strong evidence for associations between pain intensity and connectivity across four a-priori selected intrinsic brain networks. However, the replicability and consistency of these effects in independent data sets were low. SMN-SN connectivity at theta frequencies was the only brain measure to yield evidence for a correlation with pain intensity in the discovery, replication, and joint data set. This finding aligns with a recent meta-analysis showing increased hemodynamic SMN-SN connectivity in people with chronic pain relative to healthy controls [4]. Moreover, an association between SMN-SN connectivity and chronic pain appears conceptually plausible as enhanced attention to somatosensory input is a frequently observed phenomenon in people with chronic pain [25].

### Standard analysis

To contextualize our network results, we further assessed how pain intensity related to other, more commonly analyzed EEG features, i.e., frequency-band specific power and peak alpha frequency. A recent systematic review reported enhanced theta and beta power and a reduced peak alpha frequency in resting-state M/EEG recordings of people with chronic pain [20]. Our analyses provided evidence against correlations between these EEG features and chronic pain intensity. Together, these findings indicate that possible associations between chronic pain and commonly analyzed EEG features might be rather weak and inconsistent. Analyses of larger and more homogenous samples and replications in independent data sets are needed to clarify these relationships.

### Control analysis

To scrutinize the data quality and the sensitivity of our methods, we performed control analyses in which we replaced the dependent variable pain intensity by age, which has known associations with many EEG features [21, 22, 26]. These analyses yielded strong evidence for negative correlations between age and brain network connectivity at theta frequencies and between age and peak alpha frequency. In addition, the replicability and consistency of these effects were substantially higher than in the pain-related analyses. These findings confirm that the quality of the data and the sensitivity of the analysis are sufficient to detect known relationships between EEG and demographic parameters.

### Extended network analysis

Our network analysis focused on associations between pain intensity and a-priori-defined measures of brain network connectivity. In extended analyses, we investigated whether other definitions of brain network connectivity might be equally or more informative. To this end, we trained and tested ML models using an expanded feature space, which included other spatial definitions of brain networks and other approaches to computing connectivity. The extended model showed a higher prediction performance than the initial network model. Further analyses revealed that the extended model’s predictions were predominantly driven by connectivity features based on Yeo-7 intrinsic brain networks and computed via the proposed orthogonalization method for representative signals. Moreover, we confirmed that measures of brain connectivity were more informative than measures of local brain activity. Thus, this extended network model can serve as a performance benchmark and methodological inspiration for future EEG-based prediction models of pain intensity in people with chronic pain.

### Possible reasons for low replicability

The overall low replicability and consistency of effects can be explained in different ways.

First, EEG signals do not contain sufficient information about pain intensity. Considering the converging evidence for changes in cortical function in chronic pain states in animals and humans [14, 27–35], this explanation is unlikely.

Second, EEG signals contain information about pain intensity, but connectivity is not sufficiently informative. Considering evidence for a crucial role of connectivity in shaping pain from fMRI and EEG studies in humans and animal studies [16, 17, 36–38], this explanation also appears unlikely.

Third, EEG-based connectivity contains information about pain intensity, but the current approach to estimating brain network connectivity is suboptimal. In light of the multiverse of possible connectivity analyses, we deem this explanation possible. Although we derived all analytical choices from theoretical and empirical considerations, a different connectivity measure (e.g., phase-instead of amplitude-based), a different source reconstruction algorithm (e.g., minimum norm instead of beamforming), a different approach for computing band-specific signal envelopes (e.g., wavelets rather than Hilbert transform), or simply a different epoch length might have been better choices. Future studies might systematically explore the multiverse of connectivity analyses to find the most informative approach.

Fourth, the current approach to EEG connectivity is appropriate, but the data are too heterogeneous and noisy. Considering the many different types of chronic pain included in the present study, this explanation appears plausible. Different chronic pain types are differently shaped by nociceptive, neuropathic, and nociplastic components whose brain mechanisms likely differ. In the trade-off between sample size and homogeneity and in view of the available data sets, we opted for a large but heterogeneous sample of people with chronic pain. Future studies might focus on more homogenous populations. Beyond, the EEG recording conditions and clinical assessments of the present study were also heterogeneous. Thus, data quality likely differed across data sets. However, considering the lack of validated measures for EEG data quality, we did not exclude data sets based on potential data quality issues. Future studies might reduce the heterogeneity of assessments and data quality by standardizing EEG recording conditions and clinical assessments.

Fifth, while the replication data might be too heterogeneous, one could argue that the discovery data were too homogeneous. To identify robust and generalizable effects, future studies should aim for discovery data that encompass the heterogeneity of potential replication data sets. Moreover, to effectively identify effects in the presence of data heterogeneity, discovery data sets must be large. Large data sets with realistic degrees of heterogeneity may be curated by combining data from diverse origins, ideally achieved via collaborative initiatives.

### Conclusion

The present study investigated associations between brain network connectivity and pain intensity in people with chronic pain. Furthermore, we assessed the replicability and consistency of effects across seven independent data sets. Despite strong evidence in the discovery set, the replicability of effects in independent data was low. We, however, found evidence for chronic pain being related to Yeo-7 connectivity patterns, and specifically SMN-SN connectivity at theta frequencies. Conversely, our analyses yielded robust evidence against associations between chronic pain and standard EEG features. Together, these findings underscore the need to assess replicability in independent data and highlight the potential of brain connectivity patterns rather than standard EEG features to serve as biomarkers of chronic pain. Future research might investigate connectivity patterns in collaborative multi-center studies coordinated by initiatives such as ENIGMA-Chronic Pain [39, 40]. Ideally, such studies should include large pre-defined sample sizes, homogenous types of chronic pain, and standardized EEG and clinical assessments. The approach developed in this study could serve as a blueprint for such studies. In this way, future research promises novel insights into the brain mechanisms of chronic pain, aiding the development of clinically valuable biomarkers and eventually improving the individual treatment of chronic pain.

## Methods

### Data sets

We based our analyses on resting-state EEG recordings in people with chronic pain. We used two EEG data sets from our research group (*Set_Munich1*, *Set_Munich2*) and identified and acquired five external EEG data sets (*Set_Brisbane*, *Set_Otago*, *Set_Boulder*, *Set_Haifa*, and *Set_Seattle*). We restricted our analyses to eyes-closed EEG recordings as these have been shown to give rise to more robust results [41]. We based computations of EEG features in each individual on the earliest 192 clean 2 s epochs (with 50% overlap) of their EEG recording. This was done to ensure that we consistently used neural data from early phases of the recording. Moreover, by using a fixed number of epochs, we excluded any sample size biases in EEG feature estimates. We opted for a fixed number of 192 clean epochs as it represented a good tradeoff between the quality of feature estimates (increasing with the number of epochs) and the number of included participants (decreasing with the number of epochs).

To acquire external data sets, we approached research groups across the globe in a structured data acquisition campaign. To this end, we first created a list of 79 studies that involved resting-state EEG recordings in people with chronic pain. This list was mostly based on a systematic review of EEG and MEG biomarkers of chronic pain [20]. In addition, the list comprised studies that had been published in the year 2022. To decide which external research groups to contact, we condensed this list using three criteria:

- Number of EEG-sensors ≥ 32
- Publication date ≥ 2013 (i.e., not older than 10 years at the time of inquiry)
- Number of participants ≥ 20

Applying these criteria shortened the list to 18 studies, some of which shared a single data set. We then contacted the corresponding authors of the respective studies to inquire whether they would be willing to share their data. To those who did not reply within a few weeks, we sent a standardized reminder email. As a result, four external research groups contributed their data to the study. Additionally, we included a data set that was recorded with an EEG system comprising less than 32 channels (*Set_Boulder*). We included this data set because it was sent to us before we engaged in our systematic inquiries and because we intended to use it to test the replicability of findings in EEG setups with fewer channels. Thus, in total, we had seven independent resting-state EEG data sets at our disposal (overview in Table 1 of main text). A detailed description of all internal and external data sets is provided in the supplementary materials.

In addition to EEG recordings, we utilized the following clinical and demographic information:

- **pain intensity:** Average pain over a period of one day to four weeks prior to assessment, rated on an 11-point numerical rating scale ranging from no pain (0) to worst imaginable pain (10). The time periods to which the ratings refer in the individual studies can be inferred from Fig. S3.
- **age:** Age of the participant at the time of the recording
- **diagnosis:** Identifier for category of pain diagnosis: Chronic back pain, not chronic back pain

As the data originate from various research groups, they are heterogeneous. We aimed to mitigate any biases due to systematic differences in EEG recording systems and/or clinical assessments by computing z-scores within studies for all dependent and independent variables. Moreover, as several brain measures have been linked to age [21, 22, 42], we also regressed out age from both independent and dependent variables in all analyses, except those considering age as the dependent variable. As explained in more detail below, in multivariate analyses, z-scoring and confounder removal were done in a manner to prevent leakage of information from test or validation to training sets in the employed cross-validation procedures.

### Automatic preprocessing

EEG data from all sources were preprocessed in a uniform manner using an automatic preprocessing pipeline, which was initially proposed by [43] and adapted in [26] for the use of resting-state recordings. This pipeline represents a concatenation of several established functions from the Matlab-based EEGLAB toolbox [44]. It comprises the following steps: Downsampling to 250 Hz, line noise removal, bad channel rejection, re-referencing to average reference, independent component analysis and automated rejection of independent components labeled by a machine learning classifier as “muscle” or “eye” [45], bad segment rejection, and epoching into 2 s epochs with 1 s overlaps. In all preprocessing functions, we used the default parameter settings.

### Evaluating brain measures

In this study, we primarily assessed source-level brain connectivity and activity at theta (4 – 8 Hz), alpha (8 – <13 Hz), and beta (13 – 30 Hz) frequencies. The connectivity between two brain structures (brain networks, regions, or locations) was defined as the amplitude envelope correlation [19] between those brain structures’ representative signals (see below). We opted for an amplitude-based connectivity metric as it is conceptually close to fMRI-based connectivity assessments on which the intrinsic brain network concept is based [46]. The activity of a brain structure was defined as the log-transformed, absolute variance of signals (i.e., power) within that brain structure which could be explained by that brain structure’s representative signal.

In addition to network-related features, we computed the following standard, sensor-level EEG-features using the DISCOVER-EEG pipeline [42]: global, absolute signal power at theta, alpha, beta, and gamma (60 – 100 Hz) frequencies and two versions of the peak alpha frequency (local maximum and center of gravity of the power spectral density in the alpha band). Herein, the global signal power in a given band was determined by averaging the power spectral density within that band and across all sensors.

#### Source reconstruction

To reconstruct source-level brain activity, we employed Linearly Constrained Minimum Variance (LCMV) beamformers [47] implemented in the Matlab-based FieldTrip toolbox [48]. Frequency-specific array-gain LCMV spatial filters for theta, alpha, and beta frequencies were constructed based on a lead field and a frequency-specific covariance matrix. The lead field was computed by a boundary element approximation of the solution to the bioelectromagnetic forward problem for a realistically shaped, three-shell head model. The covariance matrix was estimated for each individual frequency band based on band-pass filtered EEG epochs (2 s length, 1 s overlap). To ensure a robust inversion of the covariance matrix, we employed Tikhonov regularization as implemented in FieldTrip with a regularization parameter value of 5% of the average sensor power. The fixed orientation of the lead field for every source location was chosen to maximize the variance of the spatial filter output. Source-level signals were then obtained by applying the frequency-specific LCMV operator to the band-pass filtered sensor-level time series. Spatial filters were computed for source locations corresponding to the centroids of brain parcels described by the Schaefer atlas [49]. In all analyses except the extended network analysis, we used the variant of the Schafer atlas comprising 400 parcels. In extended network analyses, we used both the variant of the Schaefer atlas comprising 400, and the variant comprising 100 parcels. Subsequent analysis steps are based on the source-level signals.

#### Representative signals

To mitigate the effects of field spread, we opted for a representative signals approach to aggregate information on the level of individual brain structures (see supplementary methods for details).

The representative signal of a brain structure is commonly defined as the first principal component (PC) across all signals associated with that brain structure. A representative signal computed in this way constitutes the solution to an optimization problem by maximizing the explained variance across all signals associated with the given brain structure. In this work, we went one step further and added orthogonality constraints to this optimization problem. Specifically, we defined the representative signal of a brain structure as the one that maximizes the explained variance across all signals of that brain structure while constraining the explained variance of signals associated with other brain structures to zero. Two versions of our method exist:

- Pairwise orthogonalization: For a given brain structure pair, an associated pair of orthogonal representative time series is estimated.
- Global orthogonalization: For a given single brain structure, one representative time series is estimated that is orthogonal to a set of time series representing the activity outside of that brain structure. The number *Nc* of time series used to represent the activity outside of the brain structure of interest is a parameter of that method.

Through simulation studies (see supplementary methods for details), we found that of the considered methods, the optimal choice is global orthogonalization with *Nc* = 3.

We, therefore, employed this variant as the standard approach for extracting representative signals. In extended network analyses, which explore different connectivity definitions, we additionally considered pairwise orthogonalization and the first PC without orthogonalization.

### Statistical analysis and machine learning

To examine associations between pain intensity and neural measures at the between-subject level, we performed both univariate and multivariate analyses. In univariate analyses, we computed correlations between pain intensity ratings and single neural measures. In multivariate analyses, we computed correlations between actual pain intensity ratings and predictions of pain intensity ratings generated by machine learning (ML) models.

We integrated information from multiple data sets using a discovery + replication and a mega-analysis approach. In the discovery + replication approach, we first assessed bidirectional correlations in a designated discovery set (Set_Munich1). Depending on the direction of these correlations, we then tested for positive or negative correlations in the six replication sets, determining both the replicability and consistency of effects. *Replicability*, commonly defined as the ability to reproduce effects using the same methods but different data, was quantified in terms of the correlation between neural measures and pain intensity in the pooled replication sets, i.e., after merging the six replication sets into one. *Consistency* was quantified in terms of a consistency score which we defined as follows:

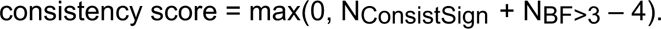

Here, NConsistSign represents the number of replication sets for which the correlation direction (positive or negative) matches that of the discovery set, and NBF>3 indicates the number of replication sets for which, additionally, correlations yield a BF > 3. By assigning equal weights to all replication sets with BF > 3, the score penalizes scenarios where replicability relies heavily on only a few or even a single replication set. The significance of a given consistency score level was estimated by simulating uncorrelated data sets (100,000 random instances) and determining the fraction of cases for which the observed score equaled or exceeded a certain level (see Fig. S4 for more details). The estimated probabilities for observing a consistency score of at least 1, 2, or 3 were p < 0.16, < 0.05, and < 0.01, respectively.

Finally, mega-analyses were performed on the joint data set, which included the discovery and all replication sets. Mega-analyses resolve the hierarchy between discovery and replication sets and are more sensitive with respect to more subtle effects present across data sets.

#### Machine learning analyses

We trained and tested ML models that relate multivariate patterns of brain connectivity across theta, alpha, and beta frequencies to the dependent variables of interest, i.e., pain intensity or age. The procedure of training and testing the ML models, which is inspired by the approaches in [17] and [16], is described in the following and visualized in Fig. S7.

First, we defined several candidate model structures. These model structures differed in the type of ML algorithm and the number of included components. Specifically, we explored two ML algorithms: principal component regression (PCR) and partial least squares regression (PLS). These algorithms were selected for their aptitude to deal with highly correlated input, as seen in EEG-based brain connectivity estimates. In basic network analyses, we explored values from one to twelve for the number of included components, which needs to be specified in both ML algorithms. Thus, basic network analyses involved 24 candidate model structures. Each model structure incorporated 18 features, corresponding to six between-network connectivity values within each of the three considered frequency bands.

We determined the best candidate model structure using cross-validation. For each candidate model structure (Fig. S7a, model selection loop), we computed a cross-validated prediction-observation correlation based on the discovery data set (Fig. S7a, inner-CV-loop) according to the following steps:

- **1. Data splitting:** Split the discovery set into training and validation sub-sets in a 9:1 ratio.
- **2. Standardization and confounder removal in training and validation sub-sets:** Standardize both model features and dependent variables by subtracting a mean and subsequently dividing by a standard deviation value. Mean and standard deviation values were estimated solely based on the training sub-set. Similarly, remove variability due to age by regressing out age from both model features and dependent variables using regression models fitted solely on the training sub-set. The described standardization and confounder removal procedure ensured that there was no information leakage from validation to training sets.
- **3. Model fitting and prediction:** Fit the model using the training set and predict observations in the validation set.
- **4. Prediction accuracy in current CV-fold:** Compute the prediction-observation correlation in the validation set.
- Repeat steps one through four for the remaining 9 data folds, then generate a new randomized data split and repeat the entire procedure 30 times.
- Finally, compute the cross-validated prediction-observation correlation as the average of the prediction-observation correlations in the 300 validation sets.

We defined the winning model structure as the one yielding the highest cross-validated prediction-observation correlation. We used the winning model structure in the final model and fitted it using the entire discovery data set.

Before assessing the model’s performance in independent data (“out-of-sample-validation”), we assessed its performance within the discovery set (“in-sample-validation”). To this end, we performed leave-one-participant-out cross-validation (LOO-CV, Fig. S7b) which involved selecting a winning model structure and fitting a final model based on all data points in the discovery set except one. The fitted model was then used to predict the target value of the omitted (test) data point. This process was iterated, leaving out a different data point in each cycle, until all data points were used as test data exactly once. As outlined above, we statistically assessed the in-sample prediction accuracy by computing Bayesian correlations across all pairs of predicted and observed target values. Importantly, in the LOO-CV loop, as in the inner-CV-loop described above, data standardization and confounder removal were performed in a manner ensuring that no information leaked from test to training sets.

To confirm the statistical significance of predictions, we also performed a permutation test (Fig. S7c). This involved repeatedly shuffling the target variable and re-calculating the cross-validated prediction-observation correlation 1000 times. Subsequently, a p-value was derived by comparing the prediction-observation correlation of the unshuffled data to the distribution of prediction-observation correlations of the shuffled data.

If the in-sample validation yielded a positive cross-validated prediction-observation correlation with BF > 3, we used the fitted model to predict observations in the standardized and confounder-corrected replication data sets. To statistically assess the out-of-sample model performance, we computed Bayesian correlations between predicted and observed target values in the replication sets.

In multivariate mega-analyses, we performed an in-sample LOO-CV (Fig. S7b) on the joint set.

#### Extended network analysis

Extended network analyses are variants of the above-described multivariate network analysis. They extend the previous approach by augmenting the space of candidate features. In addition to the two ML algorithms (PCR and PLS) and variable numbers of components (21 in the case of the extended network analysis), the new model space also encompassed multiple spatial configurations, varied techniques for generating representative time series, and different definitions of amplitude-based connectivity (see supplementary materials for details). Due to the enhanced model complexity and ensuing increased computational cost, we performed 100-fold-CV instead of the LOO-CV to assess in-sample prediction accuracies in the extended network analysis.

To investigate which types of brain connectivity features enabled the ML models’ predictions of pain intensity, we performed ablation analyses by training and testing ML models using a reduced candidate model space. Specifically, the reduced candidate model spaces excluded feature types that had been consistently selected in all folds of the in-sample CV of the “full” models. The first ablation analysis omitted features based on the Yeo-7 spatial configuration. The second ablation analysis omitted features based on either of the two proposed orthogonalization methods for representative signals.

Furthermore, we aimed to compare the information content about pain intensity of brain connectivity features with that of brain activity features. To this end, we trained and tested ML models similar to the previous approach, but substituted brain connectivity between brain structures with brain activity within brain structures as feature candidates.

## Supplementary materials

### Subgroup analysis: CBP

**Figure S1:**
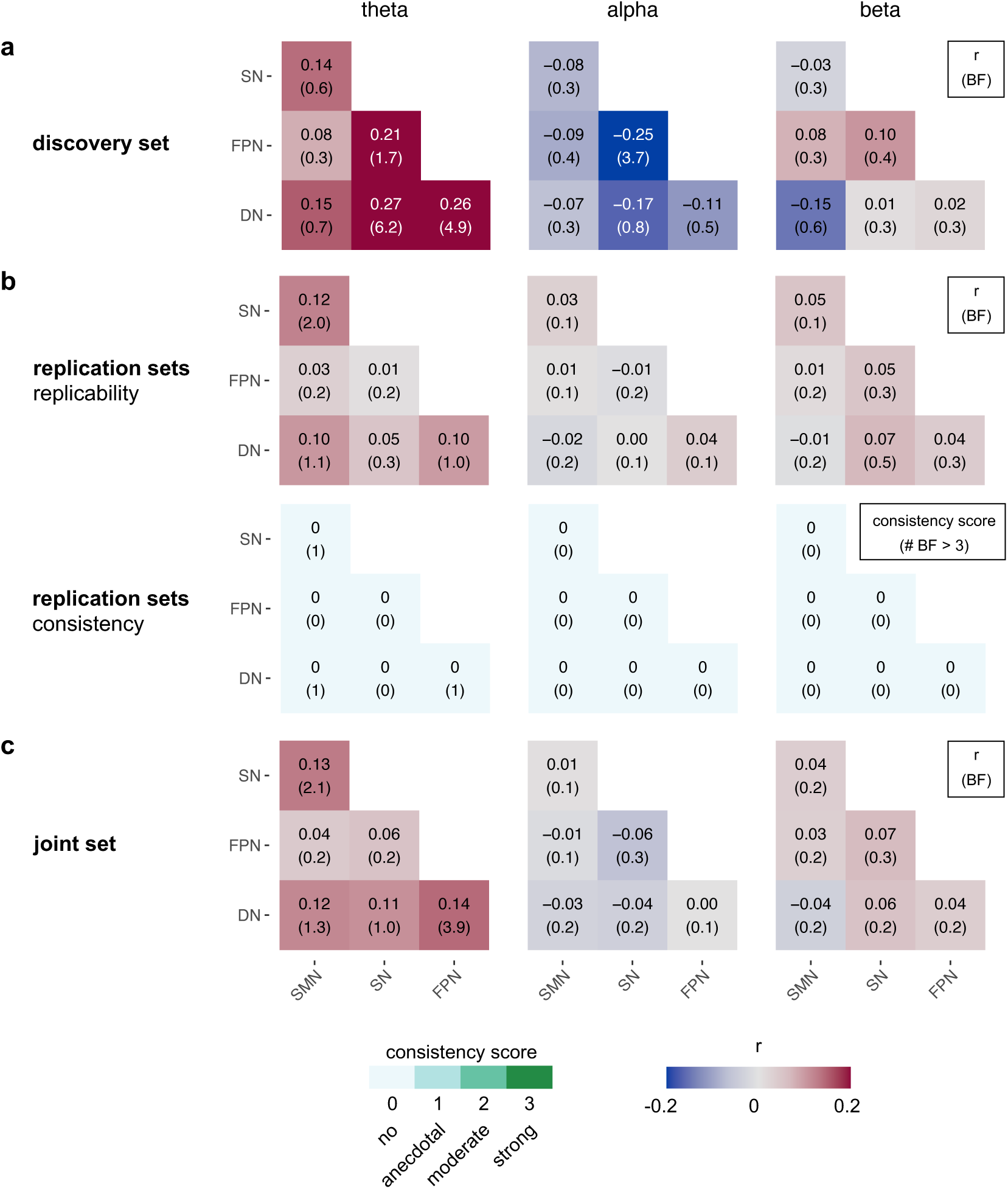
Univariate correlations between brain network connectivity at theta, alpha, and beta frequencies and pain intensity in the chronic back pain cohort. (a) Correlations between pain intensity and brain network connectivity in the discovery set. Each heatmap tile’s top number and color represent the correlation coefficient; the bottom number is the associated BF. (b) Upper panel row: Correlations in the pooled replication sets. The meanings of numbers and colors match those of panel (a). Lower panel row: Summary of correlations observed in the replication sets. The top number and color of each heatmap tile represent the consistency score. The bottom number represents the number of replication data sets that correlate with BF > 3 in the same direction as in the discovery set. (c) Correlations in the joint set. The meanings of numbers and colors match those of panel (a). SMN, somatormotor network; SN, salience network; FPN, frontoparietal network; DN, default network.

**Figure S2:**
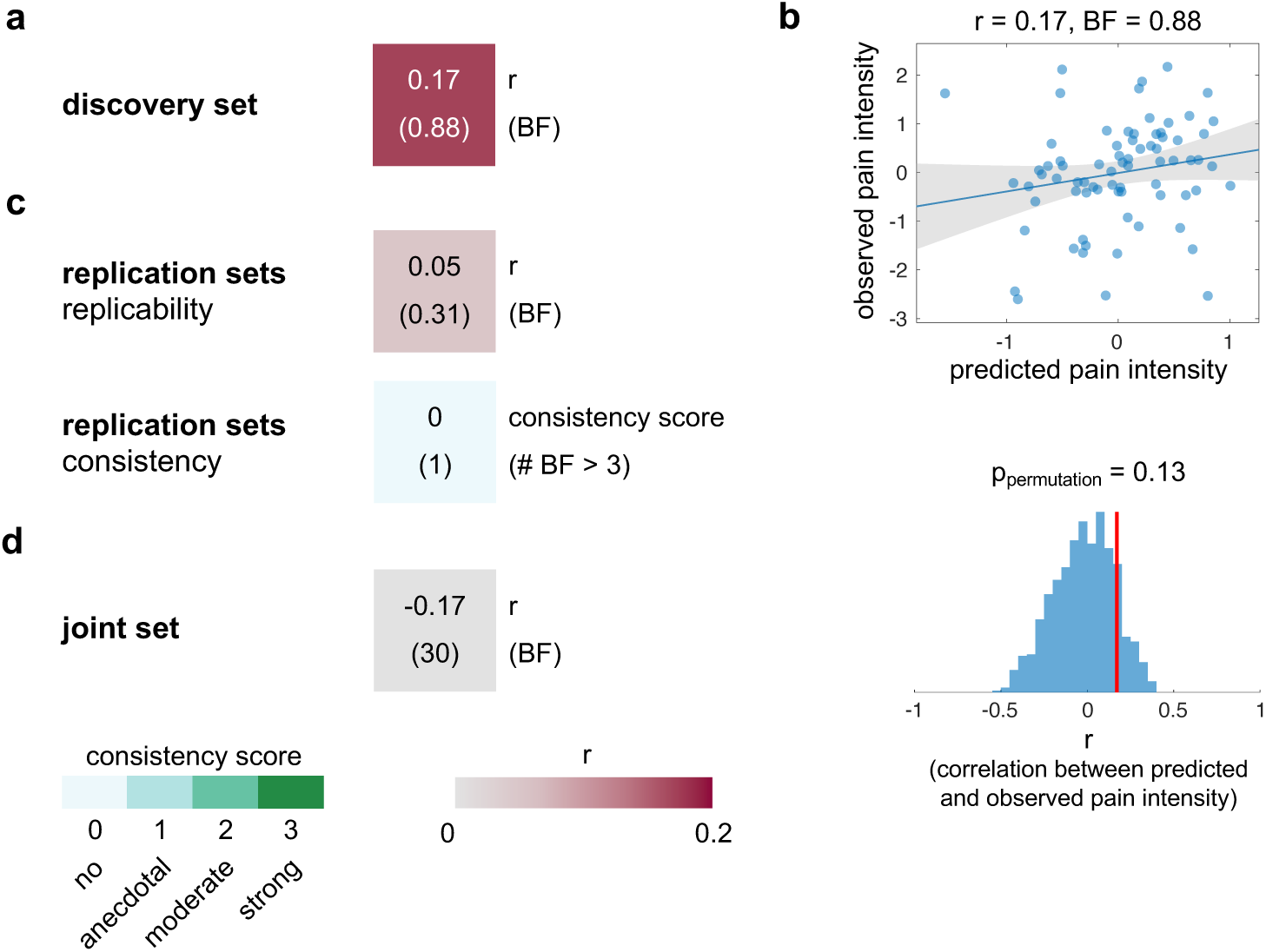
Associations between multivariate patterns of brain network connectivity and pain intensity in the chronic back pain cohort. (a) In-sample, cross-validated (LOO-CV) correlation between predicted and observed pain intensity in the discovery set. (b) Scatter plot of the cross-validated correlation between predicted and observed pain intensity and the result of a permutation test, confirming the statistical significance of predictions. (c) Upper tile: Out-of-sample correlations between predicted and observed pain intensity in the pooled replication sets. The meanings of numbers and colors are match those of panel (a). Lower tile: Summary of out-of-sample correlations observed in individual replication sets. The top number and color of each tile represent the consistency score. The bottom number represents the number of replication data sets that exhibit a positive correlation with BF > 3. (d) In-sample, cross-validated (LOO-CV) correlation between predicted and observed pain intensity in the joint set. The meanings of numbers and colors match those of panel (a).

### Data sets

*Set_Munich1* is composed of three data sets that have previously been recorded in our research group to investigate brain dysfunction in people with chronic pain. These data sets are *Set_May* [27], *Set_TaDinh* [34], and *Set_Heitmann* [50]. *Set_May* and *Set_TaDinh* have been used in combination to compare measures of brain activity, brain connectivity [34], and brain dynamics [35] (i.e., microstate analyses [51]) cross-sectionally between people with chronic pain and healthy controls. *Set_Heitmann* has been used to assess measures of brain activity and brain connectivity longitudinally in a cohort of people with chronic pain [50]. Here, we included only baseline recordings of Set_Heitmann. In all studies of *Set_Munich1*, inclusion criteria consisted of a clinical diagnosis of chronic pain, with pain persisting for at least six months and with an average pain intensity of at least four (two in the case of *Set_Heitmann2022*) on an 11-point numerical rating scale (NRS) ranging from zero (no pain) to ten (worst imaginable pain) during the four weeks prior to the assessment. People with severe diseases other than chronic pain or those taking regular benzodiazepine medication were excluded. In total, *Set_Munich1* comprised data from 127 people with chronic pain. Eight participants had to be excluded due to not meeting the minimum epoch number requirement. The analyzed cohort consisted of 74 people with chronic back pain (CBP), 13 people with chronic widespread pain (CWP), 20 people with primarily neuropathic pain (NP), and 12 people with pain of other etiologies (OTHER). All data sets were recorded using a passive electrode EEG system with 64 channels (Easycap, Herrsching, Germany) and BrainAmp MR plus amplifier (Brain Products, Munich, Germany). Previous analyses of components of *Set_Munich1* did not yield any evidence which could have biased the analyses of the present study.

*Set_Munich2* has not yet been published or analyzed, as data acquisition is still ongoing. The primary goal of this project is to assess pain medication effects on EEG-based measures of brain function. In total, data sets of 88 people with chronic pain are available, 25 of whom were excluded due to insufficient epoch numbers after preprocessing. Among the remaining n = 63 participants, 21 had CBP, two had CWP, 17 were classified as having NP, and 23 belonged to the OTHER category. To record EEG in this study, a 32-channel system with active dry electrodes (CGX-Quick32r, CGX-systems, San Diego, US) was used.

*Set_Brisbane* resulted from a study that investigated the effectiveness of several non-pharmacological, 8-week interventions for the treatment of CBP in 69 people with chronic pain. To be eligible for the study, participants had to report pain in the lower back area for more than three months with an average pain intensity of at least a 4 on an 11-point NRS ranging from 0 to 10 in the four-week period prior to assessment . Participants with severe psychiatric comorbidities were excluded from the study. After preprocessing, data from 60 participants recorded prior to interventions could be included in the present study. EEG recordings were obtained using an ANT Neuro EEGO sports system (Medical Imaging Solutions GmbH, Berlin, Germany) with 64 active scalp electrodes (Waveguard cap).

*Set_Otago* was recorded in the context of a study that investigated the efficacy of infra-slow neurofeedback training as a treatment for chronic low back pain in 60 participants [52]. Eligibility criteria were analogous to those stated for *Set_Brisbane*. For the analyses presented here, we used baseline data from 57 participants meeting the minimum epoch number condition. EEG recordings were obtained using a 64-electrode system with SynAmps-RT amplifier (Compumeics-Neuroscan, Abbotford, Australia). In the original study, no analyses relevant to this project were reported.

*Set_Boulder* is a data set for which, to date, no analyses have been published. It was recorded in the context of a larger study investigating the efficacy of pain reprocessing therapy for the treatment of chronic back pain [53]. Eligibility criteria were very similar to those described in the two previously presented data sets. *Set_Boulder* comprises data from 68 people with CBP which were recorded using a 19-channel EEG system (Evoke system, evoke neuroscience, New York, USA). No participant had to be excluded due to not meeting the minimum epoch number requirement.

*Set_Haifa* comprises recordings from 112 people with painful diabetic polyneuropathy. In the present study, we used data from 47 participants which fulfilled the minimum epoch number criterion after preprocessing. In the original study, these data were used to train a machine learning model to distinguish participants with painful from those with non-painful diabetic polyneuropathy [54]. This study was part of the larger DOLORisk [55] initiative aiming to identify risk factors for the development and maintenance of neuropathic pain. Inclusion criteria defined by this initiative were, e.g., having a diagnosis of Type 1 or Type 2 diabetes and having a clinical diagnosis of peripheral neuropathy or symptoms highly suggestive thereof. For EEG recordings, a 64-channel system with active electrodes was used (ActiCHamp, Brain Products, Munich, Germany).

Set_Seattle [56] stems from a study which investigated the relative effects of hypnotic cognitive therapy, standard cognitive therapy, hypnosis focused on pain reduction, and pain education in adults with a variety of chronic pain conditions. Here, we focus on the first five minutes of resting state EEG recordings acquired prior to the interventions. In total, data from 148 participants were available, 123 of which met our minimum epoch number criterion. Among these, 48 had CBP, the remainder of participants had chronic pain secondary to multiple sclerosis (n = 45), spinal cord injury (n = 21), or muscular dystrophy (n = 9). The inclusion criteria of the original study required potential participants to report an average pain intensity of at least four on an 11-point NRS, ranging from 0 to 10, in the past week. Moreover, participants had to report pain on at least 50% of the days in the past four weeks. Potential participants were excluded from the study if they had previously received psychological treatment or any other form of treatment akin to the treatments investigated in the study. EEG recordings were conducted using a 128-channel hydrocell net connected to a GES 300 high-density EEG acquisition system (magnetism EGI, Eugene, USA). Here, we only included data from the 116 sensors which were located in regions covered by the head model employed for source reconstruction.

### Pain intensity and age distributions

**Figure S3:**
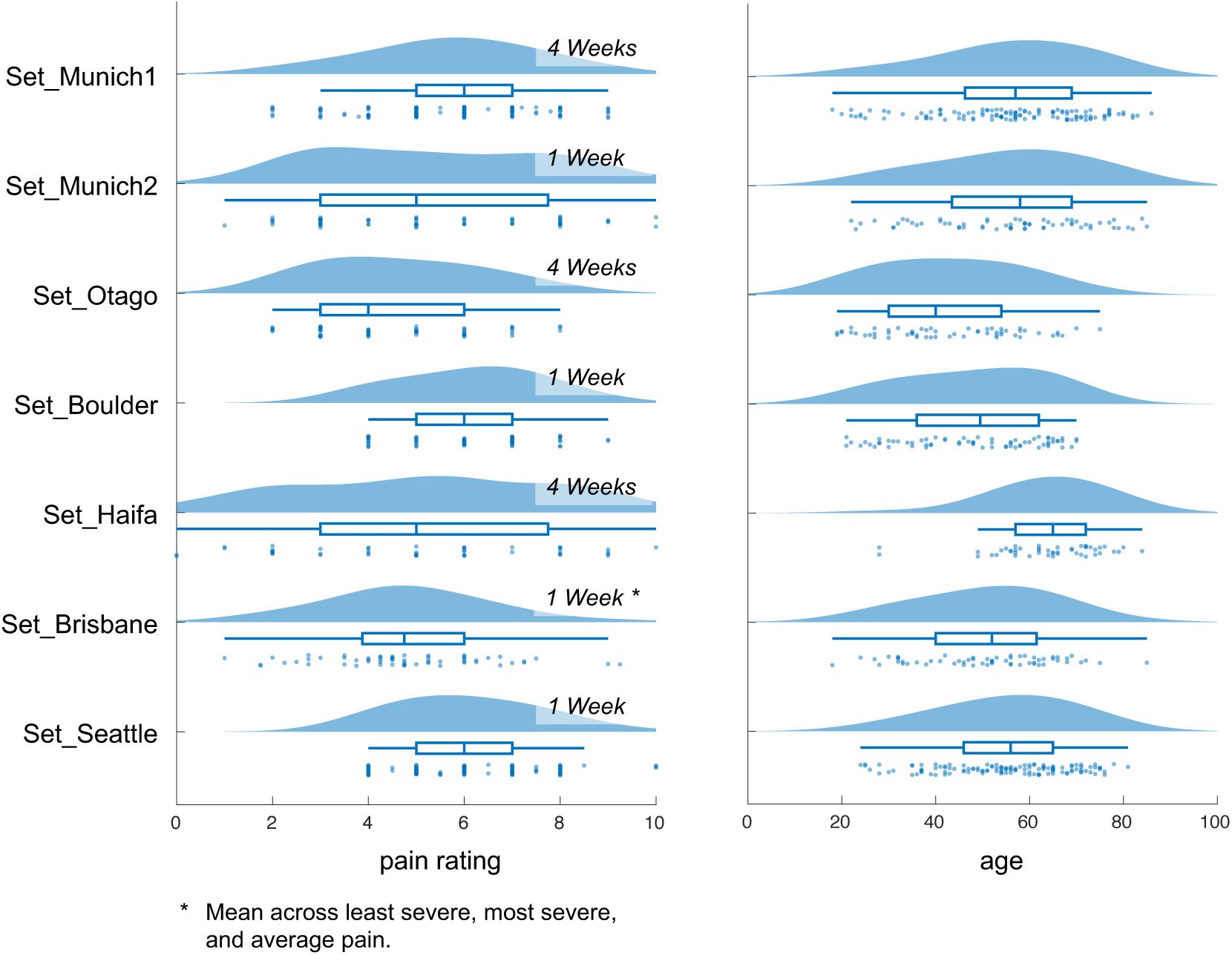
Pain intensity and age distributions. For the distributions of pain intensity, it is additionally specified for each data set to which time period prior to the assessment the ratings refer.

### Consistency score

**Figure S4:**
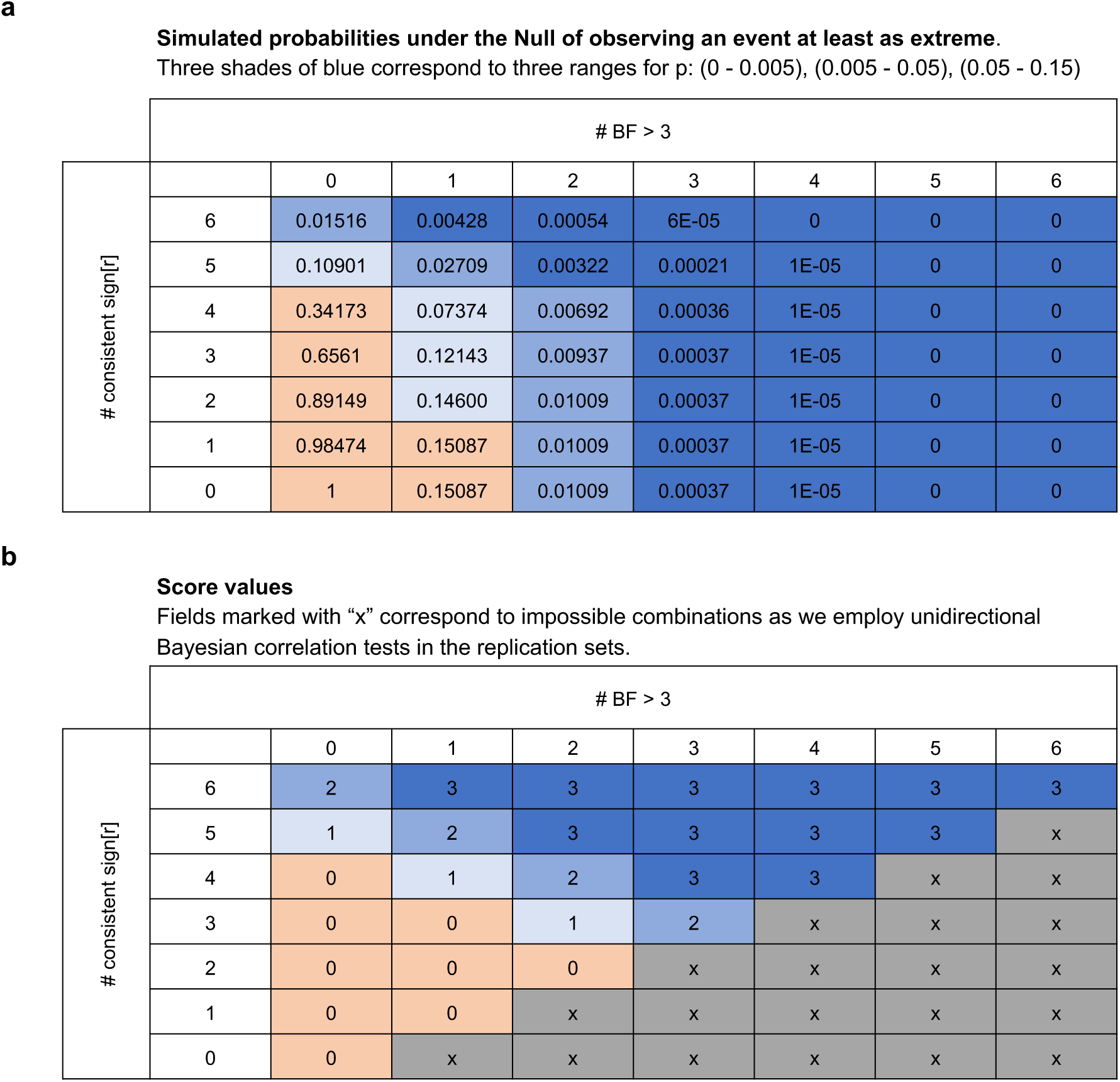
Consistency score. To assess how consistently effects of the discovery set could be replicated in independent data sets, we devised a consistency score based on the number of replication sets for which the correlation direction matched that of the discovery set (NConsistSign), and the number of replication sets for which, additionally, correlations yielded a BF > 3 (NBF>3). (a) For each combination of NConsistSign and NBF>3, we first computed the probability of observing an event at least as extreme. The results indicate that, in the relevant ranges (i.e., NConsistSign large, NBF>3 small), the probabilities have approximately the same order of magnitude for combinations where NConsistSign + NBF>3 = constant. (b) We, thus, defined a consistency score proportional to NConsistSign + NBF>3. To link the score values to quantitative statements, we estimated the probabilities of observing at least a certain score value under the Null hypothesis (see methods section of main text). For the case of six replication sets with sample sizes 63, 60, 57, 47, and 123, the probabilities under the Null hypothesis of observing a score value of a least 1, 2, or 3 are p < 0.16, < 0.05, and < 0.01, respectively. For the case of 5 replication sets with sample sizes 21, 60, 57, 68, and 48, which is relevant for the CBP subgroup analysis, the probabilities under the Null hypothesis of observing a score value of a least 1 or 2 are p < 0.08 and < 0.02, respectively. We refer to score values of 1, 2, and 3 as anecdotal, moderate, and strong evidence for consistency, respectively.

### Representative signals

**Figure S5:**
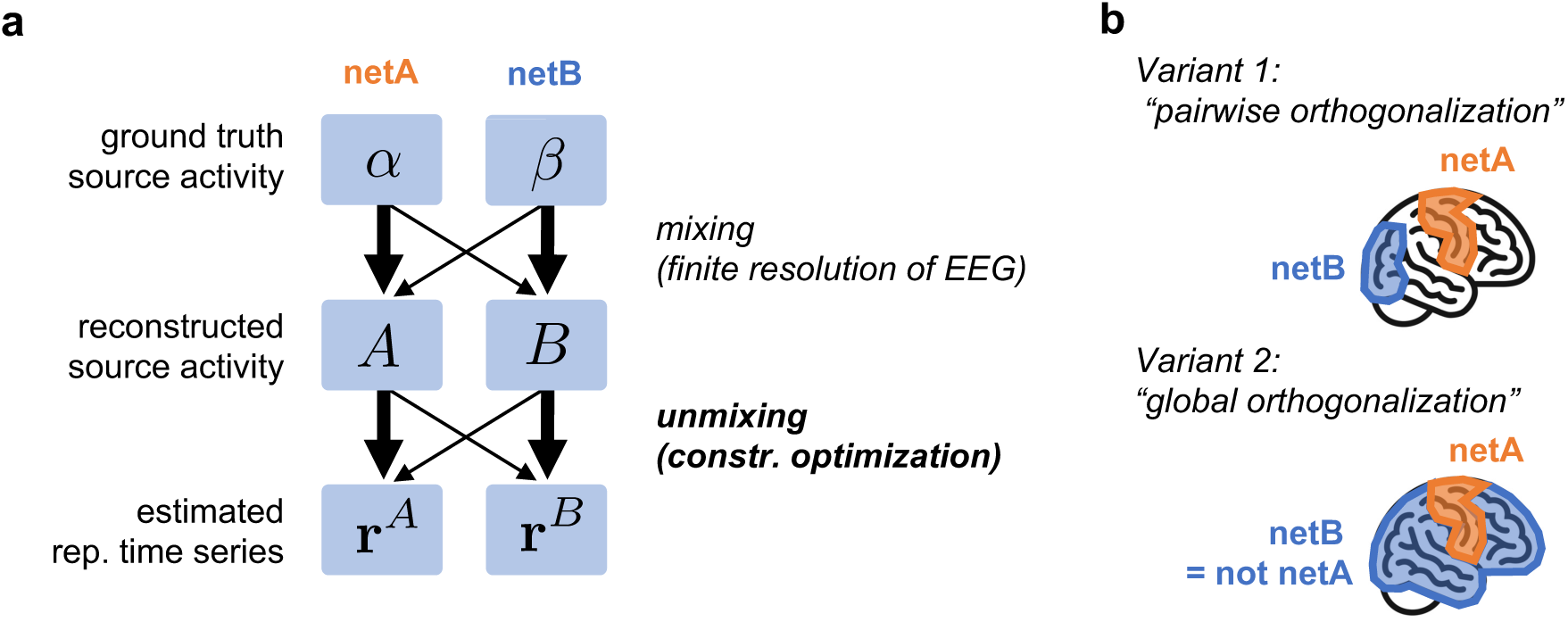
Methods for estimating representative time series. (a) Conceptual diagram of signal mixing due to imperfect source reconstruction (top half) and unmixing procedure (bottom half). (b) Illustration of two variants of the proposed method. Variant 1: For each network pair, one corresponding pair of orthogonal representative time series is determined. Variant 2: For each individual network, one representative time series is determined which is orthogonal to multiple (orthogonal) time series representing activity in all other networks.

Assessing connectivity between intrinsic brain networks using EEG is challenging due to the complex and intertwined geometries of the individual networks. A widely used method for quantifying connectivity between two smaller and simply-shaped brain structures, such as parcels or regions, is to calculate the mean of the corresponding entries of a high-resolution connectivity matrix [12, 13]. However, when applied to entire intrinsic brain networks, this approach yields connectivity values that are highly correlated across individuals (pilot assessments yielded average r-values > 0.97). This spatial indifference of connectivity values is more likely attributable to the low spatial resolution of EEG than to the true underlying network dynamics: At the low spatial resolution of EEG, a reconstructed signal at a brain source of interest is contaminated by contributions originating from surrounding sources. In particular for sources in more slender regions of a network, this means that the associated signals are contaminated by contributions from sources belonging to other networks. In other words, many signals associated with a particular network do not appropriately represent that network’s dynamics. Therefore, an alternative way to compute inter-network connectivity is to first identify signals that are believed to be sufficiently representative of the respective networks. Connectivity between networks is then determined by evaluating the connectivity between their representative signals [57].

Here, we propose a method for computing representative time series that relies on orthogonality constraints. Let’s consider networks netA and netB, for which we aim to obtain the representative time series 𝐫*^A^* and 𝐫*^B^*, respectively. These representative time series should effectively capture the ground truth activity of netA and netB. In Fig. S5a, we denote the ground truth activity of netA and netB as 𝛼 and 𝛽, respectively. However, due to the limited spatial resolution of EEG, we do not have direct access to this ground truth activity; instead, we only have access to a blurred estimate of it, which we denote as the source-reconstructed signals 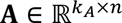 and 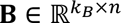 with 𝑘*_A_* and 𝑘*_B_* denoting, respectively, the number of parcels belonging to networks netA and netB and 𝑛 being the number of samples in the considered epoch. In the proposed approach, the representative time series 𝐫*^A^* and 𝐫*^B^* maximize the explained variance in 𝐀 and 𝐁, respectively, while being orthogonal. By enforcing orthogonality between 𝐫*^A^* and 𝐫*^B^*, we ensure that there is no shared portion of variance between them. Specifically, any variance in, say, 𝐫*^A^* cannot be explained by 𝐫*^B^*. As 𝐫*^B^* is constructed to be representative of the activity in netB, this orthogonalization reduces the contamination of 𝐫^!^ by the activity in netB. Formally, this approach can be expressed as the constrained optimization problem

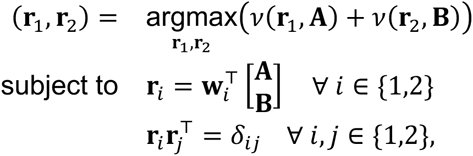

where 𝛿 is the Kronecker delta and, given that 𝐫 has unit length and the mean of the rows of 𝐗 equals zero, 𝜈(𝐫, 𝐗) is the fraction of the variance of matrix 𝐗 explained by vector 𝐫, i.e.,

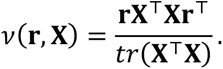

For the sake of brevity in notation, the representative time series 𝐫*^A^* and 𝐫*^B^* have been denoted by 𝐫_1_ and 𝐫_2_ respectively, in the above equation. To solve this optimization problem, a standard iterative optimization algorithm is employed, which employs information from both the local gradient and Hessian [58]. Note that, if we were to remove the orthogonality constraint 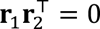, the representative time series would simply correspond to the first principal components of the data matrices 𝐀 and 𝐁.

Expanding on this idea, we can consider a variant of the method where instead of obtaining a pair of time series for each pair of networks (*“pairwise orthogonalization”*), we estimate a single time series for each network (*“global orthogonalization”*). To achieve this, we designate netA as the network of interest and define netB as all regions in the brain not contained within netA (Figure S5b). Since our focus is on obtaining a single time series only for netA, we can describe the activity of netB using multiple orthogonal components. The associated optimization problem is

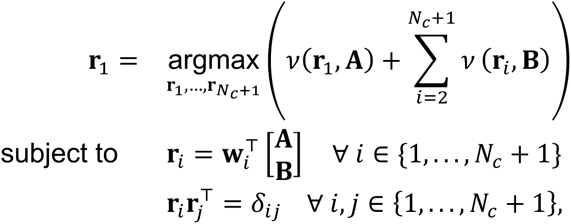

where 𝑁*_C_* denotes the number of components used to describe the activity of netB and 𝐫_1_ is the time series representative of netA. To decide which of the above-described variants should be used as the default, a simulation experiment was conducted (see below).

### Simulative assessment of representative signals method

To determine the optimal strategy for computing representative time series, a simulation experiment was conducted. In this experiment, we simulated ground truth signals at 400 locations in the brain, denoted as 𝐆 ∈ ℝ^400×*n*^. Signals differed between all 400 sources, but signals of sources belonging to the same network possessed a common component, i.e.,

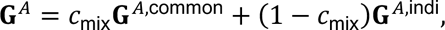

where 𝐆*^A^* refers to those rows of 𝐆 corresponding to netA. The rows of 𝐆*^A^*^,common^ are identical, while the rows of 𝐆*^A^*^,indi^ are distinct. The parameter 𝑐_mix_ controls the degree to which activity in a network is determined by the network-specific common component. To obtain the simulated source-reconstructed signals 𝐒 ∈ ℝ^400×*n*^, we applied a spatial blurring filter to the ground truth signals:

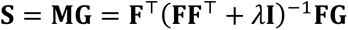

The so-called resolution matrix 𝐌 ∈ ℝ^400×400^ results from the composition of the inverse and forward models. The forward model is a linear mapping given by the lead field matrix 𝐅 and the inverse model is a linear mapping given by the minimum norm spatial filter 𝐅^T^(𝐅𝐅^T^ + 𝜆𝐈)^−1^ with 𝜆 = 𝑡𝑟(𝐅𝐅^T^)/25.

Ultimately, our objective was to compute the amplitude envelope correlation (AEC, [19]) between representative signals. For two signals 𝐫*^A^* and 𝐫*^B^*, the AEC can be defined as

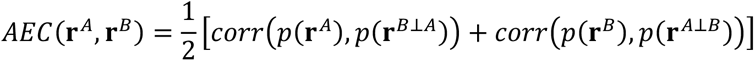

where 𝑝(𝐫) denotes the envelope of 𝐫. Further, 𝐫*^B^*^⊥*A*^ refers to the signal 𝐫*^B^* phase-orthogonalized w.r.t. 𝐫*^A^* and 𝐫*^A^*^⊥*B*^ refers to the signal 𝐫*^A^* phase-orthogonalized w.r.t. 𝐫*^B^*. Hence, there are four signals involved in the computation of the AEC: 𝐫*^A^*, 𝐫*^B^*, 𝐫*^A^*^⊥*B*^, and 𝐫*^B^*^⊥*A*^.

To quantitatively compare the different methods for estimating representative time series, we computed both the minimum and average of the following values:

- 𝜈(𝐫*^A^*, 𝐆*^A^*^,common^) = “fraction of variance of 𝐆*^A^*^,common^ explained by 𝐫*^A^*”
- 𝜈(𝐫*^A^*^⊥*B*^, 𝐆*A*,common)
- 𝜈(𝐫*B*, 𝐆*B*, common)
- 𝜈(𝐫*^B^*^⊥*A*^, 𝐆*B*,common)

Henceforth, the minimum and average of these four values are referred to as minimum and average explained variance scores, respectively.

**Figure S6:**
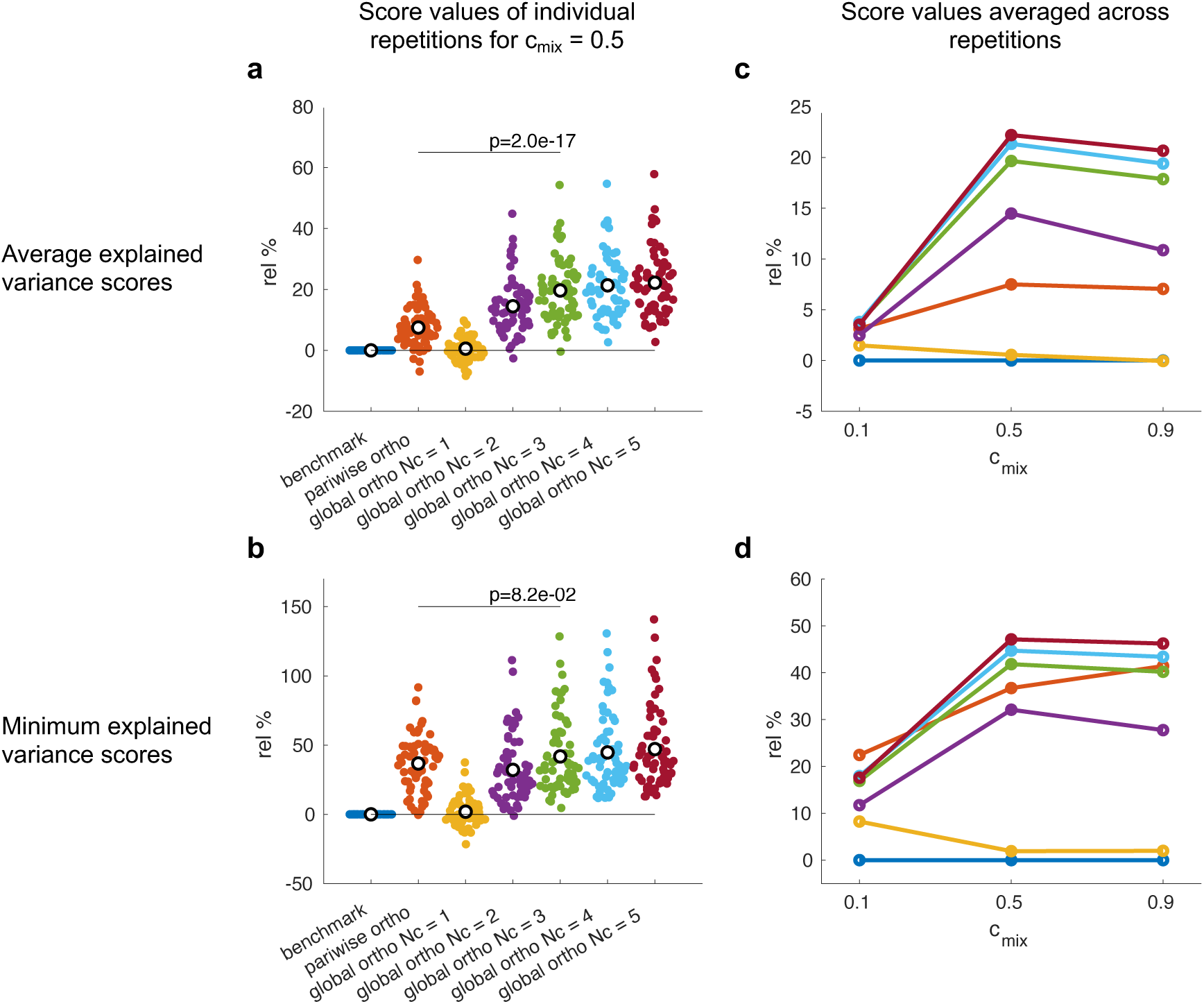
Relative explained variance scores achieved by the different methods for the Yeo-7 spatial configuration.(a), (b) Results for the case 𝑐_mix_ = 0.5 and for 64 random repetitions of the numerical experiment. Black circles indicate the mean of scores across repetitions. (c), (d) Averaged explained variance scores across repetitions for three values of the mixing parameter 𝑐_mix_.

We computed the explained variance scores for mixing parameter values 𝑐_mix_ ∈ {0.1, 0.5, 0.9}. Fig. S6 shows the explained variance scores of the different methods averaged across all network pairs and for 64 random repetitions of the simulation experiment. The values presented in the figure are relative scores, defined as the ratio between the score obtained from the method of interest and the score obtained from the standard method, i.e., standard PCA without any orthogonalization. The values in Fig. S6a and Fig. S6b correspond to the case 𝑐_mix_ = 0.5. Scores averaged across repetitions for all tested values of 𝑐_mix_ are provided in Fig. S6c and Fig. S6d.

The results demonstrate that both *pairwise* and *global* orthogonalization can significantly enhance the variance of the *common* ground truth signal that is explained by the representative time series. Among the methods tested, the best performing method is the global orthogonalization with 𝑁_1_ = 5. This method exhibits an average improvement in the average and minimum explained variance scores of roughly 20% and 50%, respectively. The variant of this method with 𝑁_1_ = 3 performs similarly well but offers the practically important advantage of being computationally less demanding. Furthermore, the pairwise orthogonalization entails a notable improvement, as well, with an increase in average and minimum explained variance scores of roughly 7% and 38%, respectively.

We conclude that the primary method for extracting representative time series should be *global orthogonalization* with 𝑁_1_ = 3.

### ML algorithm

**Figure S7:**
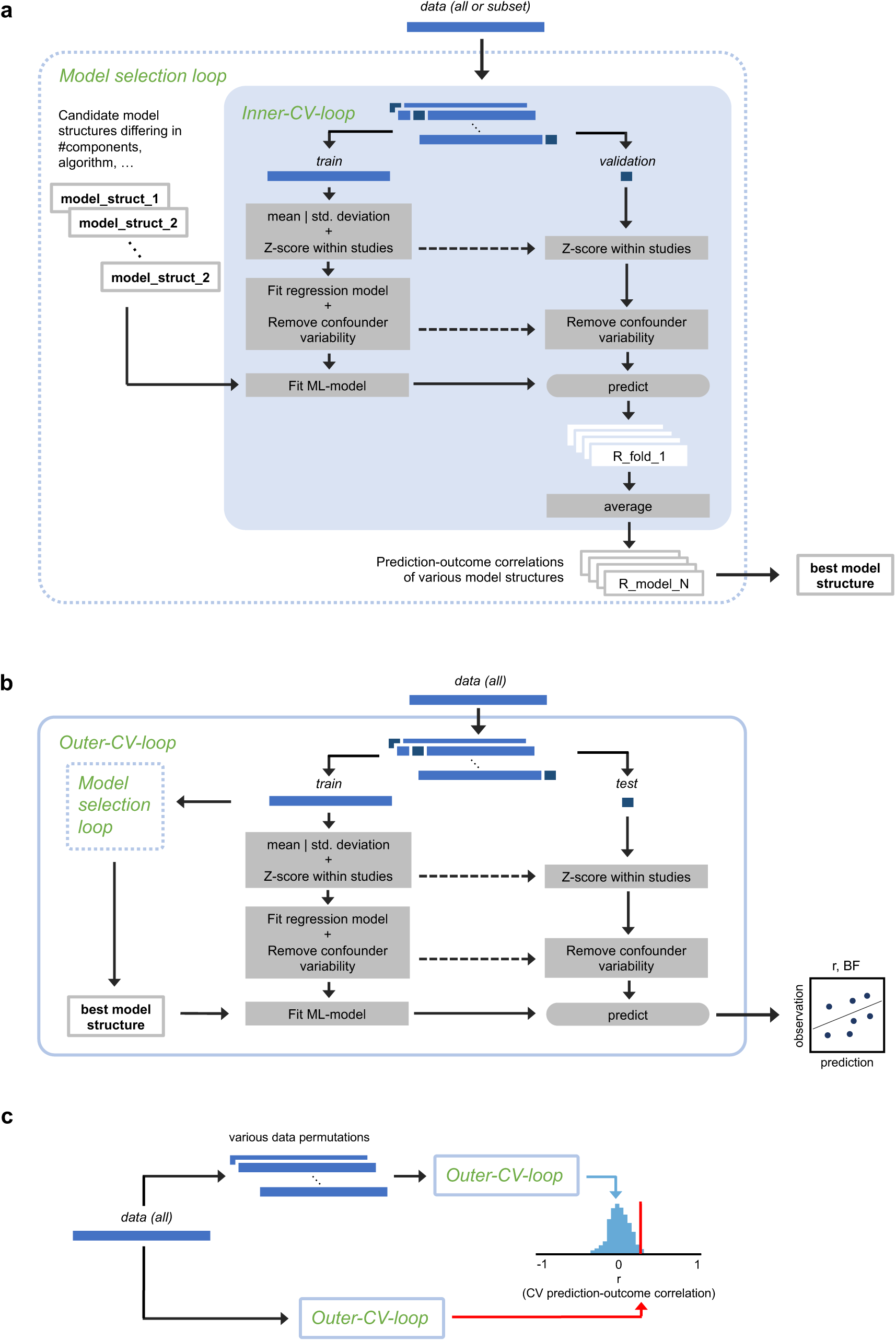
Visualization of the machine learning pipeline. (a) At the discovery stage, different model structures (i.e., machine learning algorithm/ number of included components/ spatial configuration/ connectivity metric) were assessed in a “Model selection loop” by computing cross-validated correlations between predicted and observed target values in the “Inner-CV-loop”. (b) The in-sample prediction accuracy of models resulting from the developed pipeline was estimated in an “Outer-CV-loop” using a leave-one-participant-out cross-validation procedure in the primary and a 100-fold-CV procedure in the extended network analysis. (c) To statistically confirm the prediction accuracy, we employed a permutation-based test.

### Extended network analysis

The basic network analysis considered only a single spatial configuration corresponding to four out of the seven Yeo networks. By contrast, extended network analyses involve three spatial configurations. The first configuration, labeled Yeo-7, corresponds to the complete collection of seven Yeo networks. In this configuration, the predictive model comprises 63 predictors (21 connectivity values among 7 networks in three frequency bands). The second configuration, labeled Anat-25, involves 25 anatomical brain regions which are defined by grouping the Schaefer parcels according to their anatomical labels rather than according to their network affiliation. Consequently, the predictive model comprises 900 predictors in this configuration. The third configuration, termed Schaefer-100, defines each individual parcel of the 100-parcel Schaefer atlas as a distinct spatial entity, resulting in 14,850 predictors. Visualizations of all variants of spatial configurations are provided in Fig. S8.

Moreover, the extended network analysis explored three techniques to compute representative time series. With reference to earlier explanations, these techniques are standard PCA, pairwise orthogonalization, and global orthogonalization with *Nc* = 3. Due to computational resource constraints, “pairwise orthogonalization” was not performed for the Schaefer-100 spatial configuration.

We further explored two variants of AEC definitions. The first variant, which is the one originally proposed in [19] and also employed in the earlier network analysis, involves computing correlations between logarithmically transformed squared amplitude envelopes. The second variant, which is implemented in [24], computes correlations between untransformed amplitude envelopes.

**Figure S8:**
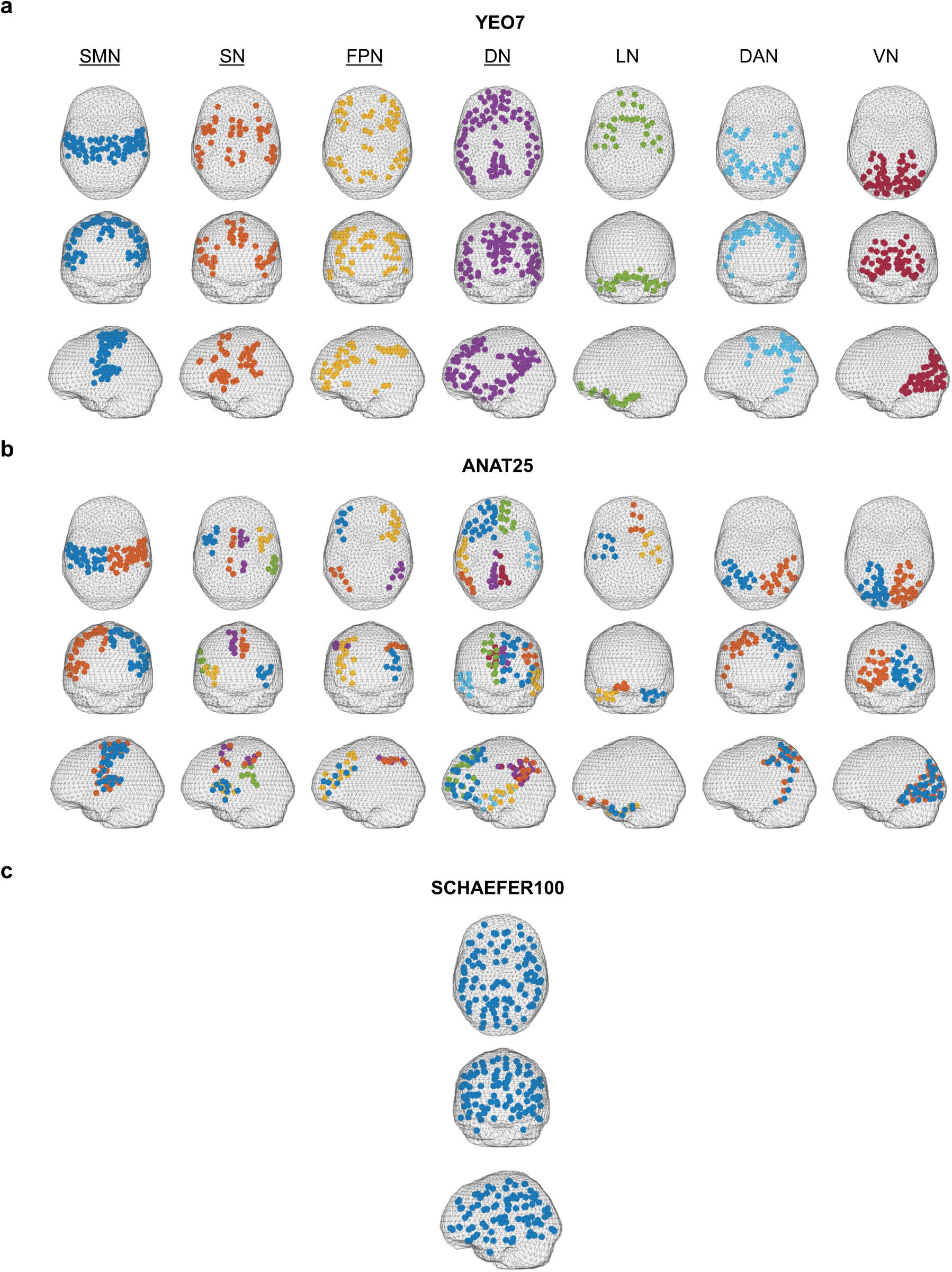
Visualization of different spatial configurations.Axial, coronal, and sagittal views of the somatormotor (SMN), the salience ventral attention (SN), the frontoparietal (FPN), the default (DN), the limbic (LN), the dorsal attention (DAN), and the visual network (VN). In primary network analyses, we focused on only four out of these seven networks (indicated by the underlined network abbreviations). In extended network analyses, we included all seven networks. (b) Axial, coronal, and sagittal views of the 25 regions of the Anat-25 spatial configuration. For visualization purposes, the Anat-25-regions are grouped by their Yeo network affiliation; that is, each Anat-25-region belongs to exactly one Yeo network. For example, two out of 25 regions correspond to the left and right halves of the SMN, another five out of 25 regions correspond to five subdomains of the SN, and so on. Different colors are used to distinguish the Anat-25-regions within each Yeo network (due to a limited palette, colors do not distinguish regions across different Yeo networks). (c) Axial, coronal, and sagittal view of the 100 parcel centroids of the Schaefer-100 spatial configuration.

## References

1. Breivik, H., et al., Survey of chronic pain in Europe: prevalence, impact on daily life, and treatment. Eur J Pain, 2006. 10(4): p. 287–333.

2. Kennedy, J., et al., Prevalence of persistent pain in the U.S. adult population: new data from the 2010 national health interview survey. J Pain, 2014. 15(10): p. 979–84.

3. Ploner, M., C. Sorg, and J. Gross, Brain Rhythms of Pain. Trends Cogn Sci, 2017. 21(2): p. 100–110.

4. Brandl, F., et al., Common and specific large-scale brain changes in major depressive disorder, anxiety disorders, and chronic pain: a transdiagnostic multimodal meta-analysis of structural and functional MRI studies. Neuropsychopharmacology, 2022.

5. Napadow, V., et al., Intrinsic brain connectivity in fibromyalgia is associated with chronic pain intensity. Arthritis Rheum, 2010. 62(8): p. 2545–55.

6. Napadow, V., et al., Decreased intrinsic brain connectivity is associated with reduced clinical pain in fibromyalgia. Arthritis Rheum, 2012. 64(7): p. 2398–403.

7. Baliki, M.N., et al., Functional Reorganization of the Default Mode Network across Chronic Pain Conditions. PLoS One, 2014. 9(9): p. e106133.

8. Hsiao, F.J., et al., Altered insula-default mode network connectivity in fibromyalgia: a resting-state magnetoencephalographic study. J Headache Pain, 2017. 18(1): p. 89.

9. Hemington, K.S., et al., Abnormal cross-network functional connectivity in chronic pain and its association with clinical symptoms. Brain Struct Funct, 2016. 221(8): p. 4203–4219.

10. Yeo, B.T., et al., The organization of the human cerebral cortex estimated by intrinsic functional connectivity. J Neurophysiol, 2011. 106(3): p. 1125–65.

11. Uddin, L.Q., B.T.T. Yeo, and R.N. Spreng, Towards a Universal Taxonomy of Macro-scale Functional Human Brain Networks. Brain Topogr, 2019. 32(6): p. 926–942.

12. Toll, R.T., et al., An Electroencephalography Connectomic Profile of Posttraumatic Stress Disorder. Am J Psychiatry, 2020. 177(3): p. 233–243.

13. Zhang, Y., et al., Identification of psychiatric disorder subtypes from functional connectivity patterns in resting-state electroencephalography. Nat Biomed Eng, 2021. 5: p. 309–323.

14. Baliki, M.N., et al., Corticostriatal functional connectivity predicts transition to chronic back pain. Nat Neurosci, 2012. 15(8): p. 1117–9.

15. Woo, C.W., et al., Distinct brain systems mediate the effects of nociceptive input and self-regulation on pain. PLoS Biol, 2015. 13(1): p. e1002036.

16. Lee, J.J., et al., A neuroimaging biomarker for sustained experimental and clinical pain. Nat Med, 2021. 27(1): p. 174–182.

17. Spisak, T., et al., Pain-free resting-state functional brain connectivity predicts individual pain sensitivity. Nat Commun, 2020. 11(1): p. 187.

18. Menon, V., Large-scale brain networks and psychopathology: a unifying triple network model. Trends Cogn Sci, 2011. 15(10): p. 483–506.

19. Hipp, J.F., et al., Large-scale cortical correlation structure of spontaneous oscillatory activity. Nat Neurosci, 2012. 15(6): p. 884–90.

20. Zebhauser, P.T., V.D. Hohn, and M. Ploner, Resting-state electroencephalography and magnetoencephalography as biomarkers of chronic pain: a systematic review. Pain, 2022.

21. Cole, J.H. and K. Franke, Predicting Age Using Neuroimaging: Innovative Brain Ageing Biomarkers. Trends Neurosci, 2017. 40(12): p. 681–690.

22. Engemann, D.A., et al., A reusable benchmark of brain-age prediction from M/EEG resting-state signals. Neuroimage, 2022. 262: p. 119521.

23. Gil Ávila, C., et al. DISCOVER-EEG: an open, automated EEG pipeline for biomarker discovery. 2023; Available from: https://osf.io/mru42/.

24. Tadel, F., et al., Brainstorm: a user-friendly application for MEG/EEG analysis. Comput Intell Neurosci, 2011. 2011: p. 879716.

25. Broadbent, P., C. Liossi, and D.E. Schoth, Attentional bias to somatosensory stimuli in chronic pain patients: a systematic review and meta-analysis. Pain, 2021. 162(2): p. 332–352.

26. Gil Ávila, C., et al., DISCOVER-EEG: an open, fully automated EEG pipeline for biomarker discovery in clinical neuroscience. bioRxiv, 2023.

27. May, E.S., et al., Prefrontal gamma oscillations reflect ongoing pain intensity in chronic back pain patients. Hum Brain Mapp, 2019.

28. Hashmi, J.A., et al., Shape shifting pain: chronification of back pain shifts brain representation from nociceptive to emotional circuits. Brain, 2013. 136(Pt 9): p. 2751–68.

29. Zebhauser, P.T., V.D. Hohn, and M. Ploner, Resting-state electroencephalography and magnetoencephalography as biomarkers of chronic pain: a systematic review. Pain, 2023. 164(6): p. 1200–1221.

30. Kuner, R. and T. Kuner, Cellular Circuits in the Brain and Their Modulation in Acute and Chronic Pain. Physiol Rev, 2021. 101(1): p. 213–258.

31. Kucyi, A. and K.D. Davis, The dynamic pain connectome. Trends Neurosci, 2015. 38(2): p. 86–95.

32. Lee, M., et al., Activation of corticostriatal circuitry relieves chronic neuropathic pain. J Neurosci, 2015. 35(13): p. 5247–59.

33. Zhou, H., et al., Inhibition of the Prefrontal Projection to the Nucleus Accumbens Enhances Pain Sensitivity and Affect. Front Cell Neurosci, 2018. 12: p. 240.

34. Ta Dinh, S., et al., Brain dysfunction in chronic pain patients assessed by resting-state electroencephalography. Pain, 2019. 160(12): p. 2751–2765.

35. May, E.S., et al., Dynamics of brain function in chronic pain patients assessed by microstate analysis of resting-state electroencephalography. Pain, 2021((May) Department of Neurology, Technical University of Munich (TUM), Germany TUM-Neuroimaging Center, Germany Center for Interdisciplinary Pain Medicine, School of Medicine, TUM, Munich, Germany).

36. Bott, F.S., et al., Local brain oscillations and interregional connectivity differentially serve sensory and expectation effects on pain. Sci Adv, 2023. 9(16): p. eadd7572.

37. Gan, Z., et al., Layer-specific pain relief pathways originating from primary motor cortex. Science, 2022. 378(6626): p. 1336–1343.

38. Stegemann, A., et al., Prefrontal engrams of long-term fear memory perpetuate pain perception. Nat Neurosci, 2023. 26(5): p. 820–829.

39. Thompson, P.M., et al., The ENIGMA Consortium: large-scale collaborative analyses of neuroimaging and genetic data. Brain Imaging Behav, 2014. 8(2): p. 153–82.

40. Quidé, Y., et al., ENIGMA-Chronic Pain: a worldwide initiative to identify brain correlates of chronic pain. PAIN, 2024. in press.

41. Duan, W., et al., Reproducibility of power spectrum, functional connectivity and network construction in resting-state EEG. Journal of Neuroscience Methods, 2021. 348.

42. Gil Avila, C., et al., DISCOVER-EEG: an open, fully automated EEG pipeline for biomarker discovery in clinical neuroscience. Sci Data, 2023. 10(1): p. 613.

43. Pernet, C.R., et al., From BIDS-Formatted EEG Data to Sensor-Space Group Results: A Fully Reproducible Workflow With EEGLAB and LIMO EEG. Front Neurosci, 2020. 14: p. 610388.

44. Delorme, A. and S. Makeig, EEGLAB: an open source toolbox for analysis of single-trial EEG dynamics including independent component analysis. Journal of Neuroscience Methods, 2004. 134(1): p. 9–21.

45. Pion-Tonachini, L., K. Kreutz-Delgado, and S. Makeig, ICLabel: An automated electroencephalographic independent component classifier, dataset, and website. Neuroimage, 2019. 198: p. 181–197.

46. Engel, A.K., et al., Intrinsic coupling modes: multiscale interactions in ongoing brain activity. Neuron, 2013. 80(4): p. 867–86.

47. Van Veen, B.D., et al., Localization of brain electrical activity via linearly constrained minimum variance spatial filtering. IEEE Trans Biomed Eng, 1997. 44(9): p. 867–80.

48. Oostenveld, R., et al., FieldTrip: Open source software for advanced analysis of MEG, EEG, and invasive electrophysiological data. Comput Intell Neurosci, 2011. 2011: p. 156869.

49. Schaefer, A., et al., Local-Global Parcellation of the Human Cerebral Cortex from Intrinsic Functional Connectivity MRI. Cereb Cortex, 2018. 28(9): p. 3095–3114.

50. Heitmann, H., et al., Longitudinal resting-state electroencephalography in chronic pain patients undergoing interdisciplinary multimodal pain therapy. PAIN, 2022.

51. Khanna, A., et al., Microstates in resting-state EEG: current status and future directions. Neurosci Biobehav Rev, 2015. 49: p. 105–13.

52. Adhia, D.B., et al., Infraslow Neurofeedback Training Alters Effective Connectivity in Individuals with Chronic Low Back Pain: A Secondary Analysis of a Pilot Randomized Placebo-Controlled Study. Brain Sci, 2022. 12(11).

53. Ashar, Y.K., et al., Effect of Pain Reprocessing Therapy vs Placebo and Usual Care for Patients With Chronic Back Pain: A Randomized Clinical Trial. JAMA Psychiatry, 2022. 79(1): p. 13–23.

54. Topaz, L.S., et al., Electroencephalography functional connectivity-A biomarker for painful polyneuropathy. Eur J Neurol, 2023. 30(1): p. 204–214.

55. Pascal, M.M.V., et al., DOLORisk: study protocol for a multi-centre observational study to understand the risk factors and determinants of neuropathic pain. Wellcome Open Res, 2018. 3: p. 63.

56. Jensen, M.P., et al., Pain-related beliefs, cognitive processes, and electroencephalography band power as predictors and mediators of the effects of psychological chronic pain interventions. Pain, 2021. 162(7): p. 2036–2050.

57. Pellegrini, F., et al., Identifying good practices for detecting inter-regional linear functional connectivity from EEG. NeuroImage, 2023. 277: p. 120218.

58. Byrd, R.H., J.C. Gilbert, and J. Nocedal, A trust region method based on interior point techniques for nonlinear programming. Mathematical programming, 2000. 89: p. 149–185.

